## Supplementary Information for "The SARS-CoV-2 nucleocapsid protein is dynamic, disordered, and phase separates with RNA"

### Sequence analysis

Disorder prediction was performed using IUPred2.0, with additional analysis and sequence parsing done with localCIDER and protfasta, respectively [1–3].

Amino acid sequence of the N protein used in simulations. Highlighted regions delineate folded domains. Underline bolded residues identify the sites of dyes for single-molecule fluorescence experiments.

```
1    MSDNGPQNQR NAPRITFGGP SDSTGSNQNG ERSGARSKQR RPQGLPNNTA
51   SWFTALTQHG KEDLKFPRGQ GVPINTNSSP DDQIGYYRRA TRRIRGGDGK
101  MKDLSPRWYF YYLGTGPEAG LPYGANKDGI IWVATEGALN TPKDHIGTRN
151  PANNAAIVLQ LPQGTTLPKG FYAEGSRGGS QASSRSSSRS RNSSRNSTPG
201  SSRGTSPARM AGNGGDAALA LLLLDRLNQL ESKMSGKGQQ QQGQTVTKKS
251  AAEASKKPRQ KRTATKAYNV TQAFGRRGPE QTQGNFGDQE LIRQGTDYKH
301  WPQIAQFAPS ASAFFGMSRI GMEVTPSGTW LTYTGAIKLD DKDPNFKDQV
351  ILLNKHIDAY KTFPPTEPKK DKKKKADETQ ALPQRQKKQQ TVTLLPAADL
401  DDFSKQLQQS MSSADSTQA
```

A complete list of constructs is presented in **Table S1**.

### Simulation Methods

#### Monte Carlo simulations

All Monte Carlo simulations were performed using the CAMPARI simulation engine and ABSINTH implicit solvent model (`abs_3.2_opls.prm`) using the monovalent ion parameters derived by Mao et al.[4]. All simulations were performed at 330 K and at 15 mM NaCl, as have been used previously in various systems [5–8]. The base keyfile used for all Monte Carlo simulations can be found at https://github.com/holehouse-lab/supportingdata/.

Simulation analysis was performed with MDTraj and camparitraj (http://ctraj.com/) [9]. For IDR only simulations, all degrees of freedom were fully sampled (backbone and sidechain dihedral angles and rigid-body positions) as is standard in CAMPARI Monte Carlo simulations [10]. For simulations of IDRs in the context of folded domains, the backbone dihedral angles of the folded domains were held fixed, while all sidechains were fully sampled, as were backbone dihedral angles for the disordered regions, as applied previously [11]. The folded state starting structures were obtained from PDB structures obtained from molecular dynamics (MD) simulations (see below for more details).

For IDR-only simulations, 30-40 independent simulations were run generating final ensembles of 40-60 K conformations. For simulations of IDRs in the context of folded domains, the number of independent simulations and the length of the simulation varied. For the NTD-RBD simulations 400 independent simulations were run using an initial molecular dynamics based sampling approach to obtain starting states for the folded domain, with 2 independent simulations per starting seed from MD simulations (see methods below) leading to a final ensemble of ~400 K conformations (24 M steps per simulation). For the RBD-LINK-dimerization construct, thirty-five independent simulations were run for a final ensemble of 32 K conformers (66 M steps per simulation). For the dimerization-CTD construct 200 independent simulations were run providing a final ensemble of 40 K conformations (66 M steps per simulation). For a complete description of simulation details see **Table S4**.

For both the NTD-RBD construct and the DIM-CTD construct, we used a sequential sampling approach in which long timescale MD simulations of the RBD in isolation performed on the Folding@home distributed computing platform were first used to generate hundreds of starting conformations [12]. Those RBD conformations were then used as starting structures for independent all-atom Monte Carlo simulations. Monte Carlo simulations were performed with the ABSINTH forcefield in which the RBD backbone dihedral angles are held fixed but the NTD is fully sampled, as are RBD sidechains. For simulations of the monomeric dimerization domain we discovered that as a monomer, the first 21 residues of the dimerization domain appear disordered, in agreement with sequence predictions (**Fig. 1A**) but in contrast to their behavior in the dimeric structure (**Fig. 1C**). As a result, we choose to also model these residues as fully disordered.

The RBD starting structure used was taken as the first chain extracted from the 6VYO PDB crystal structure, which is structurally almost identical to many of the 6YI3 NMR model shown in **Fig. 1A.** At the time that our work on this project began the 6VYO structure was the only available structure of the RBD. Irrespective, the extensive molecular dynamics (MD) simulation run prior to our Monte Carlo simulations are such that any small difference in starting structure are negated by many microseconds of simulation sampling.

To generate the monomeric starting structure of the dimerization domain, we first built a homology model of the SARS-CoV-2 dimerization dimer from the NMR structure of the SARS dimerization structure (PDB: 2JW8) using SWISS-MODEL [13,14]. We chose this strategy because at the time, no dimerization structure existed, a situation that has since resolved itself [15,16]. Nevertheless, the SARS and SARS-CoV-2 dimerization domains are essentially identical, such that this is a minor detail. As with the RBD, the application of extensive MD simulations prior to Monte Carlo simulations negates any differences in starting structure.

For RBD-link-dimerization domain simulations (316 residue systems), we opted to use a single starting seed structure for the folded domains based on the NMR and crystal-structure conformations for the RBD and dimerization domains, respectively. During these simulations, a subset of the trajectories became stuck due to long-lived interactions between the RBD and the dimerization domain, an effect likely that rose from exposed hydrophobic residues in the dimerization domain being exposed as 'folded' residues. To mitigate the impact of these unphysiological sub-ensembles, we identified trajectories in which we found contiguous simulation frames in which 25% or more of the total simulation ensemble showed unvarying interdye distance. This diagnostic identified 3 of the 31 independent replicas as being problematic, and these were discarded from our analysis. The remaining ensemble consists of 29 independent trajectories.

Excluded volume (EV) simulations were performed using the same setup, but with a modified Hamiltonian under which solvation, attractive Lennard-Jones, and polar (charge) interactions are scaled to zero, as described previously [17].

**Molecular dynamics simulations**

All molecular dynamics simulations of SARS-CoV-2 nucleoprotein were performed with Gromacs 2019 using the AMBER03 force field with explicit TIP3P solvent [18–20]. Simulations were prepared by placing the starting structure in a dodecahedron box that extends 1.0 Å beyond the protein in any dimension. The system was then solvated, and energy minimized with a steepest descents algorithm until the maximum force fell below 100 kJ/mol/nm using a step size of 0.01 nm and a cutoff distance of 1.2 nm for the neighbor list, Coulomb interactions, and van der Waals interactions. For production runs, all bonds were constrained with the LINCS algorithm and virtual sites were used to allow a 4 fs time step [21,22]. Cutoffs of 1.1 nm were used for the neighbor list with 0.9 for Coulomb and van der Waals interactions. The Verlet cutoff scheme was used for the neighbor list. The stochastic velocity rescaling (v-rescale) thermostat was used to hold the temperature at 300 K [23]. Conformations were stored every 20 ps.

The FAST algorithm was used to enhance conformational sampling and quickly explore the dominant motions of nucleoprotein [24,25]. FAST-pocket simulations were run for 6 rounds, with 10 simulations per round, where each simulation was 40 ns in length (2.4 μs aggregate simulation). The FAST-pocket ranking function favored restarting simulations from states with large pocket openings. Additionally, a similarity penalty was added to the ranking to promote conformational diversity in starting structures, as has been described previously [26]. The FAST dataset was clustered using a k-centers algorithm based on RMSD between frames using backbone heavy atoms (C, Cα, Cβ, N, O) to generate 1421 discrete states, which were then launched on the distributed computing platform Folding@home [12].

To generate large-scale ensembles of the folded domains, extensive simulations on the Folding@home platform were used. For the RBD, folding@home produced 500 μs of aggregate simulation data. For a monomeric version of dimerization domain, Folding@home produced 2.12 ms of aggregate simulation data. For each of these datasets, a final k-centers clustering was performed with the combined Folding@home and FAST data using Enspara (https://github.com/bowman-lab/enspara) [27]. This clustering was performed the same as described above and generated 200 discrete states that capture maximal diversity in the conformational ensemble of the two folded domains. These states were then used as the starting seeds for the folded domain conformations in CAMPARI simulations.

**Sequential MD/MC sampling approach**

The NTD and RBD combined are 173 residues of folded and disordered protein, while the dimerization domain and CTD combined are almost exactly the same size at 172 residues. Systems of this size raises a significant challenge for all-atom sampling. To address this we leveraged a novel approach in which we first ran long all-atom molecular dynamics simulations of folded domains alone using the Folding@Home platform and the FAST approach for enhanced conformational sampling [12,25]. From each of the trajectories of the RBD or dimerization domain, we then identified 200 conformationally distinct states based on these simulations which we used as “seeds" for the starting structures of the folded domains in our Monte Carlo simulations. Using these seeds, we reconstructed the previously missing disordered regions (NTD and CTD, respectively) and ran all-atom Monte Carlo simulations in which the disordered regions are fully sampled, the folded domain sidechains are fully sampled, but the folded domains backbone dihedral angles are held fixed. For the NTD-RBD construct we ran two replicas of each starting conformation were run, with 400 independent simulations generating a total ensemble of ~400 K conformations. For the dimerization domain we did not run independent replicas from the same starting configuration, such that 200 independent simulations were run that generated an ensemble of 200 K conformations. In parallel, we also ran simulations of the NTD and CTD in isolation, enabling an assessment of the impact of the folded domain.

**Coarse-grained Polymer Simulations**

Coarse-grained simulations were performed using the PIMMS software package [5,28]. PIMMS is a Monte Carlo lattice-based simulation engine in which each bead engages in anisotropic interactions with every adjacent lattice site. Moves used here were cluster translation/rotation moves and single-bead perturbation moves. Specifically, every simulation step, each bead in the system is sampled to move to adjacent sites in random order $50^3$ of times multiplied by a factor that reflects the length of the chain. Every 100 moves (on average) a cluster of chains is randomly selected and translated or rotated, where a cluster reflects a collection of two or more chains in direct contact. This moveset provides changes to the system that reflect physical movements expected in a dynamical system, allowing us to - for equivalently sized systems - compare the apparent dynamics of assembly, as has been done previously [29–32]. We repeated the simulations presented using a range of different movesets and, while convergence varied from set-to-set, we always observed analogous results.

All simulations were performed in a 70 x 70 x 70 lattice-site box using period boundary conditions. The results reported are averaged over the final 20% of the simulation to give average values after equivalent numbers of MC steps. The “polymer” is represented as a 61-residue polymer with either a central high-affinity binding site or not. The binder is a 2-bead species. Every simulation was run for 20 x $10^9$ Monte Carlo steps, with four independent replicas. Simulations were run with 1,2,3,4 or 5 polymers and 50, 75, 100, 125, 150, 175, 200, 250, 300, 400 binders.

To further explore the physical basis for single-chain polymer condensates we ran additional extended simulations for 60 x $10^9$ Monte Carlo with a moveset that includes the ability for clusters to move. Simulations were run using the same conditions for other simulations, with ten independent simulations for condition (**Fig. S18**).

**Extended discussion on coarse-grained simulations**

For simulations of homopolymeric polymers as shown in **Fig. 6C,D** the balance of chain-compaction and phase separation is determined in part through chain length and binder $K_d$. In our system the polymer is largely unbound in the one-phase regime (suggesting the concentration of ligand in the one-phase space is below the $K_d$) but entirely coated in the two-phase regime, consistent with highly-cooperative binding behavior. In the limit of long, multivalent polymers with multivalent binders, the sharpness of the coil-to-globule transition is such that an effective two-state description of the chain emerges, in which the chain is *either* expanded (non-phase separation-competent) OR compact (coated with binders, phase separation competent).

An alternative framework for understanding our simulations of single-polymer condensates comes from the idea of two distinct concentration (phase) boundaries - one for binder:high affinity site interaction ($c_1$), and a second boundary for “nonspecific” binder:polymer interactions ($c_2$) at a higher concentration. $c_2$ reflects the boundary observed in **Fig. 6C** that delineated the one and two-phase regimes. At global concentrations below $c_2$, (but above $c_1$) the clustering of binders at a high affinity site raises the apparent local concentration of binders above $c_2$, from the perspective of other beads on the chain. In this way, a local high affinity binding site can drive “local” phase separation of a single polymer.

### Protein expression, purification, and labeling.

**Plasmid construct design.**
SARS-CoV2 Nucleocapsid protein (NCBI Reference Sequence: YP_009724397.2) including an N term extension containing **His$_9$**-*HRV 3C protease site* –
**CATCATCACCATCATCATCATCACCAC***CTCGAAGTTCTGTTCCAAGGCCCG*ATGAGTGA
TAACGGTCCCCAGAATCAACGGAATGCGCCCAGAATCACGTTCGGCGGTCCAAGCGACAG
TACAGGTTCGAATCAGAATGGTGAACGCTCTGGGGCCCGAAGCAAACAGCGTCGTCCACA
GGGTTTGCCGAACAATACGGCTAGCTGGTTCACTGCGCTGACGCAGCACGGAAAAGAAGA
CTTAAAATTTCCGCGAGGCCAGGGGGTCCCGATTAATACTAACTCCTCCCCTGACGATCAA
ATTGGTTATTATCGTCGTGCAACCCGCCGTATCCGCGGCGGAGACGGTAAAATGAAAGATC
TGTCACCGCGCTGGTATTTTTACTACCTGGGAACAGGTCCTGAAGCAGGCTTGCCGTATGG
CGCTAACAAAGATGGCATTATCTGGGTGGCTACCGAGGGTGCCCTTAATACGCCGAAAGA
TCATATTGGAACCCGTAACCCAGCCAATAACGCAGCAATCGTACTGCAGCTGCCGCAGGG
GACAACCCTGCCGAAAGGCTTTTATGCGGAAGGGAGTCGTGGCGGCAGCCAAGCCAGCT
CCCGTAGCTCCTCGCGCTCTCGCAACTCCTCGCGGAATAGTACACCGGGTTCATCACGCG
GCACCTCGCCGGCACGCATGGCTGGCAACGGGGGGGATGCGGCTTTGGCGTTACTTTTA
CTGGATAGGCTTAACCAGTTGGAAAGTAAAATGAGCGGTAAAGGCCAGCAGCAGCAGGGT
CAGACTGTGACCAAAAAGAGCGCGGCAGAGGCGTCGAAAAAACCTAGACAAAAGCGTACT
GCGACCAAAGCCTACAATGTTACGCAGGCATTCGGCCGGCGCGGTCCGGAACAAACCCA
GGGCAACTTTGGTGACCAGGAGCTGATTCGTCAGGGAACCGATTACAAACACTGGCCACA
GATCGCGCAATTTGCCCCCTCGGCGTCAGCCTTTTTTGGTATGTCTCGCATTGGGATGGAG
GTAACCCCGTCTGGCACGTGGCTGACGTACACGGGCGCTATAAAGCTGGATGATAAAGAT
CCGAACTTCAAAGACCAGGTGATCTTACTGAACAAACATATTGACGCCTATAAAACGTTCCC
CCCTACTGAACCTAAGAAAGATAAAAAAAAAAAGGCCGATGAAACCCAAGCGCTACCACAA
CGCCAGAAAAAGCAGCAGACCGTCACCCTCCTGCCGGCAGCGGACCTCGACGATTTTTCT
AAGCAACTGCAACAAAGCATGTCAAGCGCCGATAGTACACAGGCGTAA - was cloned into the BamHI EcoRI sites in the MCS of pGEX-6P-1 vector (GE Healthcare) to express the protein product:
GST-LEVLFQGPLGS**HHHHHHHHH***LEVLFQGP*MSDNGPQNQRNAPRITFGGPSDSTGSNQ
NGERSGARSKQRRPQGLPNNTASWFTALTQHGKEDLKFPRGQGVPINTNSSPDDQIGYYRRA
TRRIRGGDGKMKDLSPRWYFYYLGTGPEAGLPYGANKDGIIWVATEGALNTPKDHIGTRNPAN
NAAIVLQLPQGTTLPKGFYAEGSRGGSQASSRSSSRSRNSSRNSTPGSSRGTSPARMAGNGG
DAALALLLLDRLNQLESKMSGKGQQQQGQTVTKKSAAEASKKPRQKRTATKAYNVTQAFGRR
GPEQTQGNFGDQELIRQGTDYKHWPQIAQFAPSASAFFGMSRIGMEVTPSGTWLTYTGAIKLD
DKDPNFKDQVILLNKHIDAYKTFPPTEPKKDKKKKADETQALPQRQKKQQTVTLLPAADLDDFS
KQLQQSMSSADSTQA. Site-directed mutagenesis was performed on the His$_9$-SARS-CoV2 Nucleocapsid pGEX vector to create the N protein constructs (**Table S1**). All cloning and

site-directed mutagenesis steps were performed by Genewiz and sequences were verified using sanger sequencing.

**Protein Expression and Purification**.

Both GST-$His_9$-SARS-CoV2 NTD FL and LINK FL Nucleocapsid variants were expressed recombinantly in BL21 Codon-plus pRIL cells (Agilent). 4L cultures were grown in LB medium containing carbenicillin (100 ug/mL) to $OD_{600}$ ~ 0.6 and induced with 0.2 mM IPTG for 12 hours at 16ºC. Harvested cells were lysed with sonication at 4ºC in lysis buffer (50 mM Tris pH 8, 500 mM NaCl, 10% glycerol, 10 mg/mL lysozyme, 5 mM BME, cOmplete™ EDTA-free Protease Inhibitor Cocktail (Roche), DNAse I (NEB), RNAse H (NEB)). The supernatant was cleared by centrifugation (37000 rpm for 1 hr) and bound to an HisTrap FF column (GE Healthcare) in buffer A (50 mM Tris pH 8, 500 mM NaCl, 10% glycerol, 20 mM imidazole, 5 mM BME). GST-$His_9$-N protein fusion was eluted with buffer B (buffer A + 500 mM imidazole) and dialyzed into cleavage buffer (50 mM Tris pH 8, 50 mM NaCl, 10% glycerol, 1 mM DTT) with HRV 3C protease, thus cleaving the GST-$His_9$-N fusion yielding FL N protein with two additional N-term residues (GlyPro). FL N protein was then bound to an SP sepharose FF column (GE Healthcare) and eluted using a gradient of 0-100% buffer B (buffer A: 50 mM Tris pH 8, 50 mM NaCl, 10% glycerol, 5 mM BME, buffer B: buffer A + 1 M NaCl) over 100 min. Purified N protein variants were analyzed using SDS-PAGE and verified by electrospray ionization mass spectrometry (LC-MS). Concentrations were determined spectroscopically in 50 mM Tris (pH 8.0), 500 mM NaCl, 10% (v/v) glycerol using an extinction coefficient = 42530 $M^{-1}$ $cm^{-1}$

GST-$His_9$-SARS-CoV2 wild-type, RBD-FL, LINK-ΔDimer, NTD-RBD, and CTD-FL Nucleocapsid variants were expressed recombinantly in Gold BL21(DE3) cells (Agilent). 4 L cultures were grown in LB medium with carbenicillin (100 ug/mL) to $OD_{600}$ ~ 0.6 and induced with 0.2 mM IPTG for 3 hours at 37ºC. Harvested cells were lysed with sonication at 4ºC in lysis buffer (listed above). The supernatant was cleared by centrifugation (37000 rpm for 1 hr) and the pellet was resuspended in 50 mM Tris pH 8, 500 mM NaCl, 10% glycerol, 6 M Urea, 5 mM BME and incubated at 4ºC for one hour. The resuspension was cleared by centrifugation (37000 rpm for 1hr) and the GST-$His_9$-N protein in the supernatant was bound to a FF HisTrap column (GE Healthcare) in buffer A (50 mM Tris pH 8, 500 mM NaCl, 10% glycerol, 20 mM imidazole, 5 mM BME) containing 6 M Urea. The column was then washed with buffer A allowing the protein to refold on the column. The GST-$His_9$-N protein fusion was then eluted with buffer B (buffer A containing 500 mM imidazole) and dialyzed into cleavage buffer (50 mM Tris pH8, 50 mM NaCl, 10% glycerol, 1 mM DTT) containing HRV 3C protease. FL N protein was then bound to an SP sepharose FF column (GE Healthcare) and eluted using a gradient of 0-100% buffer B (buffer A: 50 mM Tris pH 8, 50 mM NaCl, 10% glycerol, 5 mM BME, buffer B: buffer A + 1 M NaCl) over 100 min. Purified N protein variants were analyzed using SDS-PAGE and/or verified by electrospray ionization mass spectrometry (LC-MS). Protein concentrations of stock solutions were determined spectroscopically in 50 mM Tris (pH 8.0), 200-500 mMNaCl, 10% (v/v) glycerol using extinction coefficients of 42530 $M^{-1}$ $cm^{-1}$(FL) , 26400$M^{-1}$ $cm^{-1}$ (LINK-ΔDimer), and 25200$M^{-1}$ $cm^{-1}$ (NTD-RBD).

GST-His9-SARS-CoV2 CTD Nucleocapsid was expressed recombinantly in Gold BL21(DE3) cells (Agilent). 4 L cultures were grown in LB medium with carbenicillin (100 ug/mL) to OD600 ~ 0.6 and induced with 0.2 mM IPTG for 3 hours at 37ºC. Harvested cells were lysed with sonication at 4ºC in lysis buffer (50 mM MES pH 6, 500 mM NaCl, 10% glycerol, 5 mM BME, 10mg/mL lysozyme). The supernatant was cleared by centrifugation (37000 rpm for 1 hr) and the GST-His9-N protein in the supernatant was bound to a FF HisTrap column (GE Healthcare) in buffer A (50 mM MES pH 6, 500 mM NaCl, 20 mM imidazole, 10% glycerol, 5 mM BME). The GST-His9-N protein fusion was then eluted with buffer B (buffer A containing 500 mM imidazole) and dialyzed into cleavage buffer (A. 50 mM MES pH 6, 50 mM NaCl, 10% glycerol, 1 mM DTT) containing HRV 3C protease. FL N protein was then bound to an SP sepharose FF column (GE Healthcare) and eluted using a gradient of 0-100% buffer B (buffer A: 50 mM MES pH 6, 50 mM NaCl, 10% glycerol, 5 mM BME, buffer B: buffer A + 1 M NaCl) over 100 min. Purified N protein was analyzed using SDS-PAGE. Protein concentrations of stock solutions were determined spectroscopically in 50 mM MES (pH 6.0), 300 mM NaCl, 10% (v/v) glycerol using an extinction coefficient = 120$M^{-1}$ $cm^{-1}$

**Choice of labeling positions.**
The choice of the labeling positions has been obtained as a compromise between flanking the regions of interest and a series of different criteria that regards the biophysics of disordered proteins, the structural properties of the protein, and the physicochemical properties of the fluorophores. In particular, we have attempted to obtain an optimal spacing of the fluorophores to ensure we could make use of the whole FRET dynamic range. A separation between 60 to 70 amino acids is expected to provide a transfer efficiency of about 0.5 for a disordered region with scaling exponent close to 0.5 and 0.8-0.9 for a folded or collapsed state with a scaling exponent of 0.33 [33]. We have attempted to avoid altering amino acids that are clearly involved in structurally relevant interactions based on inspection of known structures of the folded domains. When looking for labeling positions in a folded domain, we have aimed for surface exposed residues to maximize the accessibility of the cysteine residues during labeling. We have avoided placing fluorophores adjacent to charged residues to avoid possible interactions with the charges of the fluorophores. Finally, we have attempted to limit the effects of quenching between fluorophores and aromatic residues [34,35]. Regarding this point, tryptophan residues have been identified as major quenchers of Alexa 488 and 594 and a spacing of twenty or more residues would be optimal [34,35]. Following these criteria, we have preferred not to label the NTD construct in position 50 due to the close proximity with a tryptophan residue and opted for a residue within the structured RBD. Similarly, we have opted to insert the labels within the LINKER such that mutations were not altering the net charge of the LINK sequence. Finally, for the CTD we have opted for spacing the labeling position far apart from the tryptophan residue within the folded dimerization domain, though this may not have been sufficient based on the ns-FCS observations of the CTD-FL.

**Fluorescent Dye Labeling.**
All Nucleocapsid variants were labeled with Alexa Fluor 488 maleimide (Molecular Probes) under denaturing conditions in buffer A (50 mM Tris pH8, 50 mM NaCl, 10% glycerol, 6M Urea, 1 mM DTT) at a dye/protein molar ratio of 0.7/1 for 2 hrs at room temperature. Single labeled protein was isolated via ion-exchange chromatography (Mono S 5/50 GL, GE Healthcare - protein bound in buffer A and eluted with 0-100% buffer B (buffer A + 1 M NaCl) gradient over 100 min) and UV-Vis spectroscopic analysis to identify fractions with 1:1 dye:protein labeling. Single labeled Alexa Fluor 488 maleimide labeled N protein was then subsequently labeled with Alexa Fluor 594 maleimide at a dye/protein molar ratio of 1.3/1 for 2 hrs at room temperature. Double labeled (488:594) protein was then further purified via ion-exchange chromatography (Mono S 5/50 GL, GE Healthcare - see above).

### Single Molecule Spectroscopy

**Experimental setup and procedure.**
Single-molecule fluorescence measurements were performed with a Picoquant MT200 instrument (Picoquant, Germany). For single-molecule FRET measurements, a diode laser (LDH-D-C-485, PicoQuant, Germany) was synchronized with a supercontinuum laser (SuperK Extreme, NKT Photonics, Denmark), filtered by a z582/15 band pass filter (Chroma) and pulsed at 20 MHz for pulsed interleaved excitation (PIE) [36] of labeled molecules. Emitted photons were collected with a 60x1.2 UPlanSApo Superapochromat water immersion objective (Olympus, Japan), passed through a dichroic mirror (ZT568rpc, Chroma, USA), and filtered by a 100 µm pinhole (Thorlabs, USA). Photons are counted and accumulated by a HydraHarp 400 TCSPC module (Picoquant, Germany). For FRET-FCS measurements, the same diode laser was used in continuous-wave mode to excite the donor dye. Photons emitted from the sample were collected by the objective, and scattered light was suppressed by a filter (HQ500LP, Chroma Technology) before the emitted photons passed the confocal pinhole (100 mm diameter). The emitted photons were then distributed into four channels, first by a polarizing beam splitter and then by a dichroic mirror (585DCXR, Chroma) for each polarization. Donor and acceptor emission was filtered (ET525/50m or HQ642/80m, respectively, Chroma Technology) and then focused on SPAD detectors (Excelitas, USA). The arrival time of every detected photon was recorded with a HydraHarp 400 TCSPC module (PicoQuant, Germany).

FRET experiments were performed by exciting the donor dye with a laser power of 100 µW (measured at the back aperture of the objective). For pulsed interleaved excitation experiments, the power used for exciting the acceptor dye was adjusted to match a total emission intensity after acceptor excitation to the one observed upon donor excitation (between 50 and 70 mW). Single-molecule FRET efficiency histograms were acquired from samples with protein concentrations between 50 pM and 100 pM. Trigger times for excitation pulses (repetition rate 20 MHz) and photon detection events were stored with 16 ps resolution.

For fluorescence correlation spectroscopy (FCS) experiments, acceptor-donor labeled samples with a concentration of 100 pM were excited by either the 485 nm diode laser or the supercontinuum laser at the powers indicated above. However, in the experiments on protein oligomerization, due to an increase in the fluorescence background upon addition of unlabeled protein above 1 μM, only the correlations corresponding to direct acceptor excitation (582 nm) have been considered reliable for the analysis.

For nsFCS, FRET samples of acceptor-donor labeled protein with a concentration of 100 pM were excited by the same diode laser but in continuum wavelength mode.

All measurements were performed in 50 mM Tris pH 7.32, 143 mM β-mercaptoethanol (for photoprotection), 0.001% Tween 20 (for surface passivation) and GdmCl at the reported concentrations (a residual concentration of 0.05-0.06 M GdmCl is present from dilution of the protein from the stock denatured sample). All measurements were performed in uncoated polymer coverslip cuvettes (Ibidi, Wisconsin, USA) and custom-made glass cuvette coated with PEG. Both materials outperform normal glass cuvette and contribute to reduced sticking of the protein to the surface. At low salt we observed improved protection from sticking when using the PEG coated cuvette.

Each sample was measured for at least 30 min at room temperature (295 ± 0.5 K).

**FRET efficiency histograms.**

Fluorescence bursts from individual molecules were identified by time-binning photons in bins of 1 ms and retaining the burst if the total number of photons detected after donor excitation was larger than at least 20. The exact threshold was selected based on the background contribution identified in the photon counting histograms with 1 ms binning. Transfer efficiencies for each burst were calculated according to $E=nA/(nA+nD)$, where $nD$ and $nA$ are the numbers of donor and acceptor photons, respectively. Corrections for background, acceptor direct excitation, channel crosstalk, differences in detector efficiencies, and quantum yields of the dyes were applied [37]. The labeling stoichiometry ratio $S$ was computed accordingly to $S = I_D/(\gamma_{PIE} I_A + I_D)$ where $I_D$ and $I_A$ represents the total intensities observed after donor and acceptor excitation and $\gamma_{PIE}$ provides a correction factor to account for differences in the detection efficiency and laser intensities. Bursts with stoichiometry corresponding to 1:1 donor:acceptor labeling (in contrast to donor and acceptor only populations) were selected and finally from the selected bursts a histogram of transfer efficiencies is constructed. Variations in the selection criteria for the stoichiometry ratio do not impact significantly the observed mean transfer efficiency (within experimental errors).

To estimate the mean transfer efficiency and deconvolve multiple populations (e.g for the NTD construct) from the transfer efficiency histograms, each population was approximated with a Gaussian peak function. For fitting more than one peak, the histogram was analyzed with a sum of Gaussian peak functions. For the conversion of transfer efficiency to distances, we used the value of the Förster radius for Alexa488 and Alexa594 previously determined and reported in literature, $R_0$ = 5.4 nm [38]. We further correct the value accounting for the dependence of the Förster radius on the solution refractive index. The changes in refractive index caused by increasing concentrations of GdmCl or KCl were measured with an Abbe refractometer (Bausch & Lomb, USA).

Finally, we estimated a systematic error on transfer efficiency of ± 0.03, based on the variation of transfer efficiency of the same reference samples after different calibrations of the instrument over the last two years. Standard deviation of the transfer efficiency for multiple repeats of the NTD, LINK, and CTD constructs is equal or less than ± 0.01. Since we aim for a comparison with simulations, here we consider the systematic error as the larger source of error and we propagate the corresponding effect on all the calculated distances.

**Fluorescence lifetimes and anisotropies analysis.**
A quantitative interpretation of this transfer efficiency in terms of distance distribution requires the investigation of protein dynamics. A first method to assess whether the transfer efficiency reports about a rigid distance (e.g. structure formation or persistent interaction with the RBD) or is the result of a dynamic average across multiple conformations is the comparison of transfer efficiency and fluorescence lifetime.The interdependence of these two factors is expected to be linear if the protein conformations are identical on both timescales (nanoseconds as detected by the fluorescence lifetime, milliseconds as computed from the number of photons in each burst). Alternatively, protein dynamics give rise to a departure from the linear relation and an analytical limit can be computed for configurations rearranging much faster than the burst duration (see SI). The dependence of the fluorescence lifetimes on transfer efficiencies determined for each burst was compared with the behavior expected for fixed distances and for a chain sampling a broad distribution of distances. For a fixed distance, *R*, the mean donor lifetime in the presence of acceptor is given by $t_D(R) = t_{D0}(1 - E(R))$, where $t_D$ is the lifetime in the absence of acceptor, and $E(R) = 1/(1 + R^6/R_0^6)$. For a chain with a dye-to-dye distance distribution *P*(*R*), the donor lifetime is $t_D = \int tI(t)dt / \int I(t)dt$, where $I(t) = I_0\, P(R)\, Exp[-t/tD(R)]\, dR$ is the time-resolved fluorescence emission intensity following donor excitation. A similar calculation can be carried out for describing the acceptor lifetime delay given by $(t_A(R) - t_{A0})/t_{D0}$ [39]. Donor and acceptor lifetimes at different concentrations of GdmCl were analyzed by fitting subpopulation-specific time-correlated photon counting histograms after donor and acceptor excitation, respectively, using a tail fit. Errors associated with the tail fit are estimated by varying the “tail” region that undergoes the fitting procedure and computing mean and standard deviation of the fit results. In computing the average of multiple measurements, errors of the single dataset are propagated accordingly.

Multiparameter detection allows also excluding possible artifacts, such as insufficient rotational averaging of the fluorophores or quenching of the dyes. Subpopulation-specific anisotropies were determined for both donor and acceptor of all three constructs for NTD, LINK, and CTD, and values were found to vary between 0.1 and 0.2 for the donor and between 0.1 and 0.2 for the acceptor, sufficiently low to assume as a good approximation for the orientational factor $\kappa^2$ = 2/3.

**Fluorescence Correlation Spectroscopy (FCS) analysis.**

In order to determine changes in the hydrodynamic radius ($R_h$) of the protein, FCS correlations were analyzed assuming 3D diffusion of the molecule across a three-dimensional Gaussian profile of the confocal volume [40]. For 1 diffusing species, and in the absence of photophysical transitions in the time scale of the lag times analyzed, this formalism amounts to the following time autocorrelation function $g(\tau) = 1 + \frac{1}{N}(1+\frac{\tau}{\tau_D})^{-1}(1+\frac{\tau}{\alpha^2\tau_D})^{-1/2}$, where $N$ is the average number of molecules in the confocal volume, $\tau_D$ is the diffusion time along the xy plane, $\alpha$ is the eccentricity of the three dimensional Gaussian observational volume. $\tau_D = \omega_{xy}^2 / 4D$, where $D$ is the 3D translational diffusion coefficient and $\omega_{xy}$ is the radius from the center of the laser beam at which the light intensity decreases $e^2$ times from its maximum value at the center. $\alpha = \omega_z/\omega_{xy}$.

Additionally, in order to account for contributions of the photophysics of the fluorophore to the correlation observed in the μs timescale, we added two triplet terms multiplying the diffusion correlation term (see for example work by Krichevsky [41]). The overall equation that we fit to the FCS traces is then $g(\tau) = 1 + (g_{diff}(\tau) - 1)(1 + c_{T1}Exp[-\frac{\tau}{\tau_{T1}}])(1 + c_{T2}Exp[-\frac{\tau}{\tau_{T2}}])$ where $\tau_{T1}$, $\tau_{T2}$, $c_{T1}$, and $c_{T1}$, denotes the characteristic times and amplitudes of the contributions of two triplet states to $g(\tau)$. Parameters $\tau_{diff}$, $\tau_{T1}$, $\tau_{T2}$, $c_{T1}$, $c_{T2}$ and $N$ were fitted by least square nonlinear regression analysis for each concentration of unlabeled protein tested (**Fig. S14 A-B**), while $\alpha$ was fixed at a value of 6 determined independently from analysis of fluorescence intensity profiles of fluorescent nanobeads.

Making use of the definition of $\tau_{diff}$ and the Stokes-Einstein equation, we have, for each concentration of unlabeled protein ($\tau_{diff} / \tau_{diff0}$) = ($R_h / R_{h0}$), where $\tau_{diff0}$ and $R_{h0}$ are the diffusion time and hydrodynamic radius in the absence of unlabeled protein, respectively. Error bars in **Fig. S14 B** are the standard errors of $R_h / R_{h0}$ estimated from propagation of the standard errors across multiple measurements of the diffusion times obtained from the fit.

**Nanosecond Fluorescence Correlation Spectroscopy.**

Autocorrelation curves of acceptor and donor channels and cross-correlation curves between acceptor and donor channels were calculated with the methods described previously [42,43]. All samples have been measured at a concentration of 100 pM and bursts with a transfer efficiency between 0.3 and 0.8 have been selected to eliminate the contribution of donor only to the correlation amplitude. Finally, the correlation was computed over a time window of 5 μs and characteristics timescales were extracted according to:

$$g_{ij}(\tau) = 1 + \frac{1}{N}(1 - c_{AB}Exp[-(\tau - \tau_0)/\tau_{AB}])(1 + c_{CD}Exp[-(\tau - \tau_0)/\tau_{CD}])(1 + c_T Exp[-(\tau - \tau_0)/\tau_T]) \quad \text{Eq S1}$$

where N is the mean number of molecules in the confocal volume and $i$ and $j$ indicate the type of signal (either from the Acceptor or Donor channels). The three multiplicative terms describe the contribution to amplitude and timescale of photon antibunching (AB), chain dynamics (CD), and triplet blinking of the dyes (T). $\tau_{CD}$ is then converted in the reconfiguration time of the interdye distance $\tau_r$ correcting for the filtering effect of FRET as described previously [44]. An additional multiplicative CD term has been added only for the donor-donor correlations to describe the fast decay observed at very short time. Such a decay is not found in the correlations of other disordered proteins measured on the instrument and we associate the fast decay with the rotational motion of the overall protein. A fit to this fast decay is about 2 ns.

**Polymer models of distance distributions.**

Conversion of mean transfer efficiencies for fast rearranging ensembles requires the assumption of a distribution of distances. Here, we compared the results of two distinct polymer models: the Gaussian model and a Self-Avoiding Walk (SAW) model that accounts for changes in the excluded volume [45]. This second model has been shown to provide a better description of chain distribution and scaling exponent when compared to distance distributions from MD simulations [46]. Importantly, both models rely only on one single fitting parameter, the root mean square interdye distance $r = \langle R^2 \rangle^{1/2}$ for the Gaussian chain and the scaling exponent $\boldsymbol{\nu}$ for the SAW model.

Estimates of these parameters are obtained by numerically solving:

$$\langle E \rangle = \int_0^{l_c} P(R)\, E(R)\, dr \quad \text{Eq. S2}$$

where $R$ is the interdye distance, $l_c$ is the contour length of the chain, $P(r)$ represents the chosen distribution, and $E(R)$ is the Förster equation for the dependence of transfer efficiency on distance R and Förster radius:

$$E(R) = \frac{R_0^6}{R_0^6 + R^6}\,. \quad \text{Eq. S3}$$

The Gaussian chain distribution is given by:

$$P_{FJC}(R, r) = 4\pi R^2 \left(\frac{3}{2\pi r^2}\right)^{3/2} exp\left(-\frac{3R^2}{2r^2}\right) \quad \text{Eq. S4}$$

The SAW model can be expressed as:

$$P_{SAW}(R,v) = A_1 \frac{4\pi}{b_0 N^v} \left(\frac{R}{b_0 N^v}\right)^{2+g} Exp\left[- A_2 \left(\frac{R}{b_0 N^v}\right)^{\delta}\right] \quad \text{Eq. S5}$$

where $A_1 = \frac{\delta}{4\pi} \frac{\Gamma[5+g/\delta]^{\frac{3+g}{2}}}{\Gamma[3+g/\delta]^{\frac{5+g}{2}}}$ , $A_2 = \left(\frac{\Gamma[5+g/\delta]}{\Gamma[3+g/\delta]}\right)^{\frac{\delta}{2}}$, $g = \frac{(\gamma-1)}{v}$ , $\delta = \frac{1}{(1-v)}$ , $\gamma =$ 1.1615, $\Gamma$ is the Euler Gamma Function, $b_0$ = 0.55 nm is an empirical prefactor [46] , $N$ is the number of residues between the fluorophores, and $\boldsymbol{v}$ is the scaling exponent.

Finally, when converting the distance from transfer efficiencies, to account for the length of dye linkers and compare the experimental data with simulations, the root-mean-squared interdye distance *r* was rescaled according to $r_{m,n} = |m-n|^{0.5} l_{\text{dye}} |m-n+2\, l_{\text{dye}}|^{0.5}$ with $l_{\text{dye}} = 4.5$ [47,48]. Finally, the persistence length is computed using the Gaussian conversion $r^2 = 2\, l_p l_c$ [49].

**Binding of denaturant and folding.**

As in previous works [50–52], we model the chain expansion with the denaturant in terms of a simple binding model:

$$r(c) = r_0 \left(1 + \rho \frac{Kc}{1+Kc}\right) \quad \text{Eq. S6}$$

Where $r_0$ is the mean square interdye distance at zero denaturant, $\rho$ is a term the captures the extent of chain expansion with the denaturant compared to $r_0$, and the $K$ is the binding constant, and $c$ is the concentration of denaturant.

In presence of folded domains, we can imagine the folding/unfolding of the domains can affect the overall size of the chain because of an increase or decrease of excluded volume due to the surrounding folded domains (which screen part of the available conformations) or because of the folding or unfolding of elements in the region between the fluorophores. To account for this effect, as in the case of the NTD, we weighed the effect of denaturant on the chain for the fraction folded $f_f$ and unfolded $f_u$ accordingly to:

$$r(c) = (r_{0f} f_f + r_{0u} f_u) \left(1 + \rho \frac{Kc}{1+Kc}\right) \quad \text{Eq. S7}$$

where $r_{0f}$ and $r_{0u}$ are the root mean square interdye distance in presence of folded or unfolded domains in native buffer,

$$f_f = \frac{Exp[-m(c-c_{1/2})/RT]}{1+Exp[-m(c-c_{1/2})/RT]} \quad \text{Eq. S8}$$

and $f_u = 1 - f_f$, where $c_{1/2}$ is the midpoint concentration and $m$ the denaturant m value, representing the dependence of free energy on denaturant concentration. The stability parameter $\Delta G_0$ can be computed as $\Delta G_0 = m\, c_{1/2}$ .

**Folding of RBD domain.** While characterizing the NTD denaturant dependence, we discovered a plateau at transfer efficiencies between 1 and 2 M GdmCl, which we interpret as the contribution of the coexistence of folding and unfolding conformations (**Eq. S7)**. To test whether this corresponds to the actual range of the folding transition, we designed, expressed, and labeled a construct with dyes in position 68 and 172, which directly monitors the folding of this domain. Single-molecule FRET measurements reveal up to three distinct populations (**Fig. S6**). One is abundant at high GdmCl concentration and disappears at low GdmCl concentrations and therefore we assign it as an unfolded state. Another one is only transiently populated between 1 and 2 M GdmCl and we assign it as an intermediate folding state. A third one, with a higher transfer efficiency compatible with the distance expected from the known RBD structure, is stabilized below 2 M GdmCl and, therefore, is assigned as the folded configuration. In absence of evident differences in brightness between these three species, the relative area of each state represents the fraction of the corresponding population. We use a three-state model where the fraction of each state can be computed from the partition function of the system, leading to:

$$f_u = \frac{1}{1+K^{u-i}+K^{i-f}} \; ; \; f_i = \frac{1}{1+(K^{u-i})^{-1}+K^{i-f}} \; ; \quad f_f = \frac{1}{1+(K^{u-i})^{-1}(K^{i-f})^{-1}+(K^{i-f})^{-1}} \qquad \text{Eq. S9}$$

where $K^{u-i}$ and $K^{i-f}$ are

$$K^{u-i} = Exp[-m^{u-i}(c-c_{1/2}^{u-i})/RT] \quad ; \quad K^{i-f} = Exp[-m^{i-f}(c-c_{1/2}^{i-f})/RT] \qquad \text{Eq. S10}$$

Fitted values to the model are reported in **Table S2.** Importantly, the observed values confirm in large measure the inferred stability measured via the NTD. The small discrepancy in the overall stability observed (**Fig. S9**) can either be assigned to the complicated decoupling of folding and chain expansion when observing the transition from the perspective of the NTD or by the “local” nature of the RBD unfolding probed by the NTD.

**Salt dependence of NTD, LINK, and CTD conformations.**

In addition to studying the conformations under native buffer conditions, we investigate how salt affects the conformations of the three disordered regions. We started by testing the effects of electrostatic interactions on the NTD conformational ensemble. Moving from buffer conditions and increasing concentration of KCl, we observed a small but noticeable shift toward lower transfer efficiencies, which represents an expansion of the NTD due to screening of electrostatic interactions. This can be rationalized in terms of the polyampholyte theory of Higgs and Joanny [50,53] (see **Table S3**), where the increasing concentration of ions screens the interaction between oppositely charged residues (see **Fig. S11**).

We then analyzed for comparison the LINK FL construct. Interestingly, we find a negligible effect of salt screening on the root mean square distance of the low transfer efficiency population as measured by FRET (see **Fig. S11**). Predictions of the Higgs & Joanny theory (see SI) for the content of negative and positive charges within the LINK construct indicates a variation of interdye distance dimension that is comparable with the measurement error. It has to be noted that in this case the excluded volume term in the Higgs and Joanny theory will empirically account not only for the excluded volume of the amino acids in the chain, but also for the excluded volume occupied by the two folded domains.

To better understand the weak dependence on salt (and denaturant) of the dimensions LINK FL and the occurrence of two populations at low salt screening, we further investigated a truncated version of the same protein, the LINK-ΔDimer construct. First of all, we observe a sharp collapse as a function of GdmCl (**Fig. S8**), which starkly contrasts with the weak change of the LINK-FL. This strongly implies an effect of the two domains in modulating the dimensions of the LINK. We then asked whether such modulation in a low denaturant regime contains a strong electrostatic component. To separate the effect of structural destabilization and electrostatic attraction in disordered proteins, we chose to use Urea. When comparing the conformation in the two denaturants, we clearly observed that Urea maintains the LINK-ΔDimer in a more compact configuration and by addition of 0.5 M KCl we can recover the expansion observed in GdmCl (**Fig. S10**). For comparison no change is observed when studying the NTD FL under the same conditions (**Fig. S10**). These observations for the LINK-ΔDimer mimic what was previously observed in the case of the Cold Shock Protein from *Thermotoga Maritima* [50] and confirms a strong electrostatic contribution in controlling the dimensions of the LINK region in absence of DIMER and CTD domains. It is reasonable to assume that similar electrostatic interactions are at play also in the full-length protein and are at the origin of the coexistence of two populations in low ionic strength solutions.

Finally, we test if the addition of salt can provide similar effects than those obtained by GdmCl on the conformations of the CTD: interestingly, we do not observe any significant variation either in transfer efficiency (**Fig. S11**), suggesting that the broadening of the population observed for the CTD does not originate exclusively from electrostatic interactions. However, when comparing the denaturing effect of GdmCl and Urea on the CTD-FL we observe more compact conformations of the chain in GdmCl.

**Polymer model of electrostatic interactions.**

The disordered regions of the N protein are enriched in positive and negative charges. To provide a term of comparison in the interpretation of protein conformations as function of salt concentration, we use the polymer theory for polyampholyte solutions developed by Higgs and Joanny [50,53], which has been shown previously to capture quantitatively the conformational changes of unstructured proteins. Briefly, the root mean square interdye distance is equal $r = N^{0.5} l_0 \alpha$ where $N$ is the number of monomers in the disordered region, $l_0$ is the length of elementary segment (here 0.36 nm) and $\alpha$ is the ratio between $l$ and $l_0$, with $l$ being a rescaled segment that accounts for excluded volume and electrostatic interactions.

$\alpha$ is computed according to the equation proposed by Higgs and Joanny [50,53]:

$$\alpha^5 - \alpha^3 = \frac{4}{3} \left(\frac{3}{2\pi}\right)^{1.5} N^{0.5} v^* \qquad \text{Eq. S13}$$

where $v^*$ is an effective excluded volume given by the sum of three terms:

$$\text{v}^* b^3 = \text{v}\ b^3 + \frac{4\pi l_B (f-g)^2}{k^2} - \frac{\pi l_B^2 (f-g)^2}{k} \quad \text{Eq. S14}$$

Here, v is the excluded volume (accounting for physical excluded volume and positive and attractive interactions that are not due to electrostatics), $f$ and $g$ are the fraction of positive and negative residue respectively for considered segment of the protein, $k$ is the Debye screening length, and $l_b$ is the Bjerrum length.

Importantly, when accounting for the fraction of negative charges, we also account for the contribution of the -2 net charge of each dye at pH 7.3.

**Testing protein oligomerization.**

NativePAGE experiments were performed to verify that purified recombinantly expressed SARS-CoV-2 N protein is capable of forming dimers and oligomers, in analogy to SARS-CoV N protein, and as shown in more recent work for SARS-CoV-2 [13,54,55]. Indeed, NativePAGE experiments reveal the existence of multiple bands (**Fig. S14 C-D**). However, since the lowest band in the NativePAGE corresponds to an apparent molecular weight of ~70-80 kDa, we wanted to verify the oligomeric state of this band.

To test whether the apparent mass is due to a slow mobility of the protein because of its high positive charge, we performed crosslinking experiments. These experiments confirm the formation of dimers, tetramers, and high oligomeric species, as a function of protein concentration above 500 nM (**Fig. S14 E-F**). These oligomeric species are in equilibrium with the monomer, the smallest species on the denaturing SDS PAGE (which has the expected molecular weight of ~45 kDa). It has to be noted that, because of the slow reactivity of the crosslinking agent (see Methods below), the crosslinking experiments do not represent the population of monomeric and oligomeric species at equilibrium. However, the comparison between the NativePAGE and the crosslinking experiments supports the fact that the smallest band in the NativePAGE is indeed the monomer protein.

We finally turned to Fluorescence Correlation Spectroscopy (FCS) to test whether labeled protein can form dimers. We measured the CTD construct that carries one labeling position at the end of the oligomerization domain. When increasing the concentration of unlabeled protein, we observe a systematic increase in the hydrodynamic radius when compared to the hydrodynamic radius under native conditions (**Fig S14 A-B**). This suggests that the labeled protein can form higher oligomeric species in a concentration regime comparable to the one observed in NativePAGE and SDS PAGE experiments and that at 100 pM (the concentration used in single-molecule experiments), no oligomer is formed. Caution must be used in the interpretation of the oligomeric bound species observed in FCS experiments, since labeling mutation may have affected the affinity of the dimerization domain. Future experiments will address the role of mutation on dimerization. Finally, all experiments have been performed at two different time points, after 1 hour and after 24 hours of incubation of the labeled sample with unlabeled protein to test any kinetic effect on the measured value. No significant difference has been observed.

Taken together, NativePAGE crosslinking experiments verify that in smFRET and FCS experiments we are in fact monitoring the behavior of the monomeric SARS-CoV-2 N protein.

***Protein Crosslinking Methods.*** 50 mM disuccinimidyl suberate (DSS) (Thermo Scientific) stock solution was prepared (10 mg into 540 uL of anhydrous DMSO (Sigma)). All protein samples were prepared in 20 mM NaPi pH 7.4 (with and without 200 mM NaCl) at the following concentrations: 0.1, 0.5, 1, 5, 10 and 20uM. DSS stock solution was added to each sample to a final concentration of 1.25 mM. Samples were incubated for 1 hour at room temperature. Samples were then quenched to a final concentration of 200 mM Tris pH 7.4 and allowed to incubate for 15 minutes. Crosslinked proteins were then analyzed using SDS PAGE and Coomassie staining.

***NativePAGE Methods.*** All protein samples were prepared in 20 mM NaPi pH 7.4 (with and without 200 mM NaCl) at the following concentrations: 0.05, 0.1, 0.5, 1, 5, 10 and 20 μM. Samples were subjected to NativePAGE (Invitrogen) and protein mobility was analyzed with Coomassie staining.

**Turbidity measurements.**

Development of turbidity in solutions of N protein and poly(rU) was followed through measurements of absorbance at 340 nm in a microvolume spectrophotometer (NanoDrop, Thermo, USA). Mixtures were prepared in 500 μl plastic reaction tubes by adding 4 μl protein solution into 3 μl of poly(rU) and absorbance was recorded 45 s – 75 s after mixing. Working solutions were kept at room temperature during experiments.

Reaction media was 50 mM Tris, pH 7.5 (HCl), 0.002 % v/v Tween20, and NaCl as indicated in *Results*.

poly(rU) (Midland Certified Reagent Company, TX, USA, lot number 011805) was reconstituted into this media from stocks dissolved in RNAse free water. According to the manufacturer, the size of poly(rU) molecules is mostly less than 250 nucleotides (nt.) and longer than 200 nt..

Protein stocks (in 50 mM Tris pH 8.0, 500 mM NaCl, 10% v/v glycerol) were buffer exchanged into the desired buffer through size exclusion chromatography in Zeba Spin 7 k MWCO desalting columns (Thermo, USA). poly(rU) concentrations in working dilutions were assessed through the absorbance at 260 nm employing an extinction coefficient of 9.4 $mM^{-1}$ $cm^{-1}$ [56]. Protein concentrations were assessed through the absorbance at 280 nm employing an extinction coefficient of 42.53 $mM^{-1}$ $cm^{-1}$, computed according to the method proposed by Pace *et al.* [57].

The limiting concentrations of nucleic acid across which an increase in turbidity was detected were estimated through interpolation of the data. To this end, an empirical equation, describing the trends observed at all concentrations, was fitted to the data and then was solved to extract the poly(rU) concentrations at which turbidity reaches a limit value above the background signal. We used a limiting absorbance value of 0.005 units (340 nm, 1 mm path length). We found that an appropriate function for this end is an exponential of a Gaussian distribution function $F(x)$:

$$F(x) = A(1 - Exp[-\beta\gamma(x)]) \qquad \text{Eq. S11}$$

where

$$\gamma(x) = \frac{1}{(2\pi)^{0.5}\sigma} Exp[-(x-\mu)^2/2\sigma^2] \qquad \text{Eq. S12}$$

where *x* denotes poly(rU) concentration and *A*, $\beta$, *σ* and *μ* are parameters fitted through weighted minimum least squares for each protein concentration (solid lines in **Fig. 5 A-B** and limiting value points in panels **C-D**). To characterize the observed global trends of turbidity, as a function of both RNA and protein concentration, we determined approximate functional forms of the dependence on protein concentration of the individually fitted parameters (*A*(*p*), $\beta$(*p*), *σ*(*p*) and *μ*(*p*), where *p* is protein concentration). The observed dependencies were increasing linearly for *μ*(*p*) and quadratic for $\beta$(*p*) and *σ*(*p*). *A* was the worst defined parameter and thus displayed the least clear trend. For the results in absence of added salt we employed an increasing power function with exponent as a fitting parameter (best fit value was < 1), whereas for the results in presence of 50 mM NaCl the trend of *A*(*p*) was better described by a decreasing exponential function.

We thus used the functional forms *A*(*p*), $\beta$(*p*), *σ*(*p*) and *μ*(*p*) to construct a global function dependent on both protein and RNA concentration. Global fitting of this equation to the whole set of turbidity titration curves provided the turbidity contour plots shown in **Fig. 5 C-D** (solid lines). Contour lines were computed at 1, 10, 20, 50 and 100 times the limiting value employed ($A_{340\ nm,1mm}$ = 0.005).

**Table S1. N protein constructs**

| N protein constructs | Start Position | End Position | Labeling Positions |
|---|---|---|---|
| Wildtype | 1 | 419 | - |
| NTD FL | 1 | 419 | M1C-R68C |
| NTD-RBD | 1 | 173 | M1C-R68C |
| RBD FL | 1 | 419 | R68C-Y172C |
| LINK FL | 1 | 419 | Y172C-T245C |
| LINK-ΔDimer | 1 | 247 | Y172C-T245C |
| CTD FL | 1 | 419 | F363C-A419C |
| CTD | 363 | 419 | F363C-A419C |

**Table S2. Fit parameters to denaturant binding models.**

| | ρ | K ($M^{-1}$) | $R_0$ (Å) | m (kcal $mol^{-1}$ $M^{-1}$) | $c_{1/2}$ (M) |
|---|---|---|---|---|---|
| **NTD-FL (2 state)** | 1.4 ± 0.3 | 0.34 ± 0.04 | 47 ± 2 (fixed $r_{0f}$) | 4.0 ± 0.8 | 1.3± 0.2 |
| | | | 34 ± 3 ($r_{0u}$) | | |
| **NTD-RBD** | 1.5± 0.2 | 0.30± 0.03 | 46 ± 2 (fixed $r_{0f}$) | 5.7 ± 1.5 | 1.50 ± 0.09 |
| | | | 36 ± 2 ($r_{0u}$) | | |
| **NTD-FL (3 state)** | 0.97 ± 0.09 | 0.34 ± 0.04 | 47 ± 2 (fixed $r_{0f}$) | Fixed based on RBD fit | Fixed based on RBD fit |
| | | | 39.8 ± 0.9 ($r_{0u}$) | | |
| **NTD-RBD (3 state)** | 1.28 ± 0.09 | 0.32 ± 0.03 | 47 ± 2 (fixed $r_{0f}$) | Fixed based on RBD fit | Fixed based on RBD fit |
| | | | 38.3 ± 0.6 ($r_{0u}$) | | |
| **RBD-FL** | - | - | - | 6.6 ± 0.5 ($m^{U-I}$) | 1.68 ± 0.02 ($c_{1/2}^{U-I}$) |
| | | | | 8.1 ± 0.5 ($m^{I-F}$) | 1.64 ± 0.02 ($c_{1/2}^{I-F}$) |
| **LINK-FL** | 0.9 ± 0.3 | 0.09 ± 0.04 | 55 ± 2 (fixed) | - | - |
| **LINK-ΔDimer** | 0.98 ± 0.07 | 0.64 ± 0.14 | 42± 2 (fixed) | - | - |
| | 0.9 (fixed based on LINK-FL) | 0.18 ± 0.04 | 49.0 ± 1.4 | - | - |
| **CTD-FL** | 0.54 ± 0.05 | 0.23 ± 0.04 | 51 ± 2 | | |
| **CTD** | 0.62 ± 0.07 | 0.42 ± 0.11 | 47.1 ± 1.4 (fixed) | | |

**Table S3. Fit parameters of Higgs & Joanny theory**

| | v |
|---|---|
| **NTD-FL** | 3.47 ± 0.05 |
| **LINK-FL** | 4.2 ± 0.2 |
| **CTD-FL** | 7.5 ± 0.5 |

**Table S3. Scaling exponents**

| | $\nu_{\text{SAW}}$ | $\nu_{\text{simulation}}$ |
|---|---|---|
| **NTD-FL** | 0.510 ± 0.009 | 0.52 |
| **NTD-RBD** | 0.500 ± 0.009 | |
| **LINK-FL** | 0.530 ± 0.008 | 0.58 |
| **LINK-ΔDIMER** | 0.464 ± 0.009 | |
| **CTD-FL** | 0.542 ± 0.007 | 0.49 |
| **CTD** | 0.534 ± 0.008 | |

**Table S4. All-atom simulation summary**

| System | No. sims | Total steps per sim (M). | Prod. steps per sim.(M) | Config. output | Ensemble size |
|---|---|---|---|---|---|
| **NTD-RBD** | 400 | 24 | 20 | 20,000 | 399,000 |
| **RBD-LINK-DIM** | 29 | 66 | 60 | 20,000 | 82,113 |
| **DIM-CTD** | 200 | 24 | 20 | 20,000 | 200,000 |
| **NTD** | 40 | 71 | 66 | 30,000 | 64,000 |
| **LINK** | 30 | 101 | 80 | 30,000 | 66,660 |
| **CTD** | 40 | 71 | 66 | 30,000 | 64,000 |

**Table S5. Interdye distances**

| | $R_{Gauss}$(Å) | $R_{SAW}$(Å) | $R_{simulation}$(Å) |
|---|---|---|---|
| **NTD-FL** | 48 ± 2 | 47 ± 2 | 53 |
| **NTD-RBD** | 46 ± 2 | 45 ± 2 | |
| **LINK-FL low E** | 55± 2 | 54 ± 2 | 59 |
| **LINK-FL high E** | 42 ± 2 | 45 ± 2 | 59 |
| **LINK-ΔDIMER** | 40 ± 2 | 40 ± 2 | |
| **CTD-FL** | 51 ± 2 | 48.7 ± 1.4 | 46 |
| **CTD** | 49 ± 2 | 47.1 ± 1.4 | |

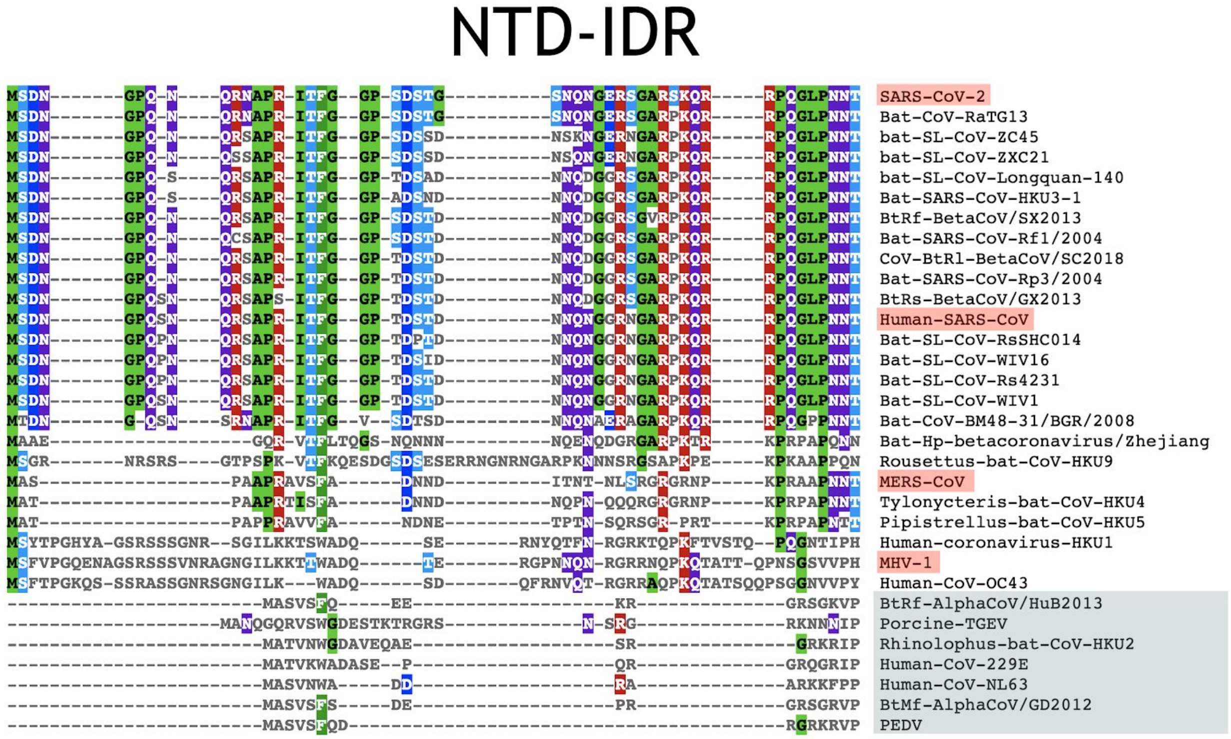


**Fig. S1. Sequence alignment of the coronavirus N-terminal domain (NTD)**

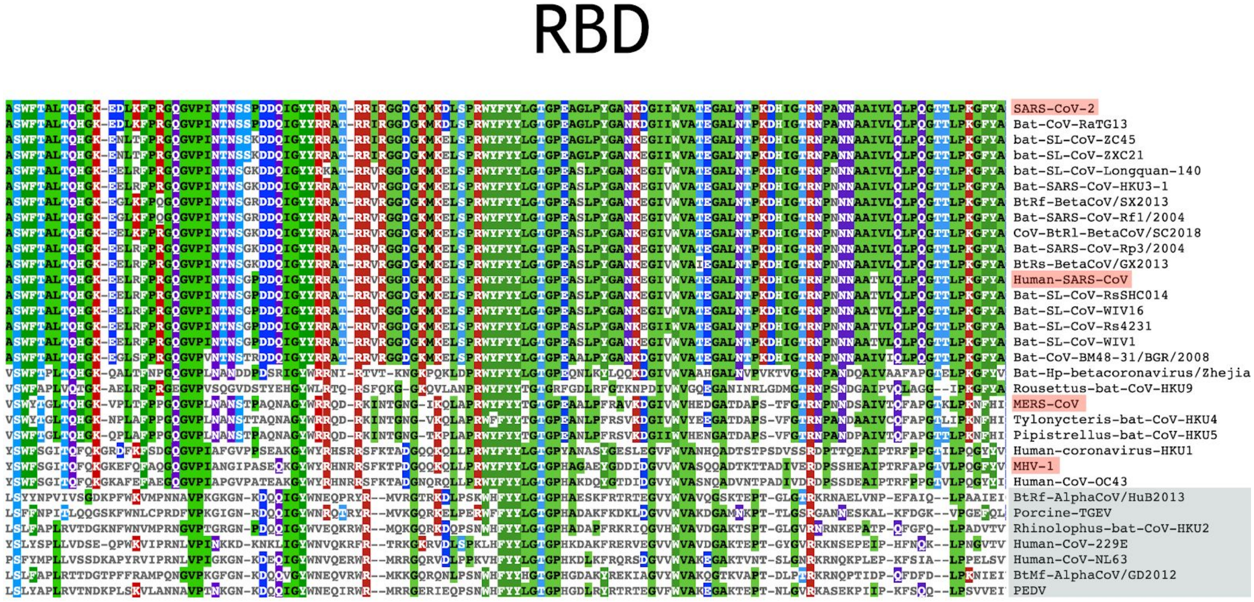


**Fig. S2. Sequence alignment of the coronavirus RNA binding domain (RBD)**

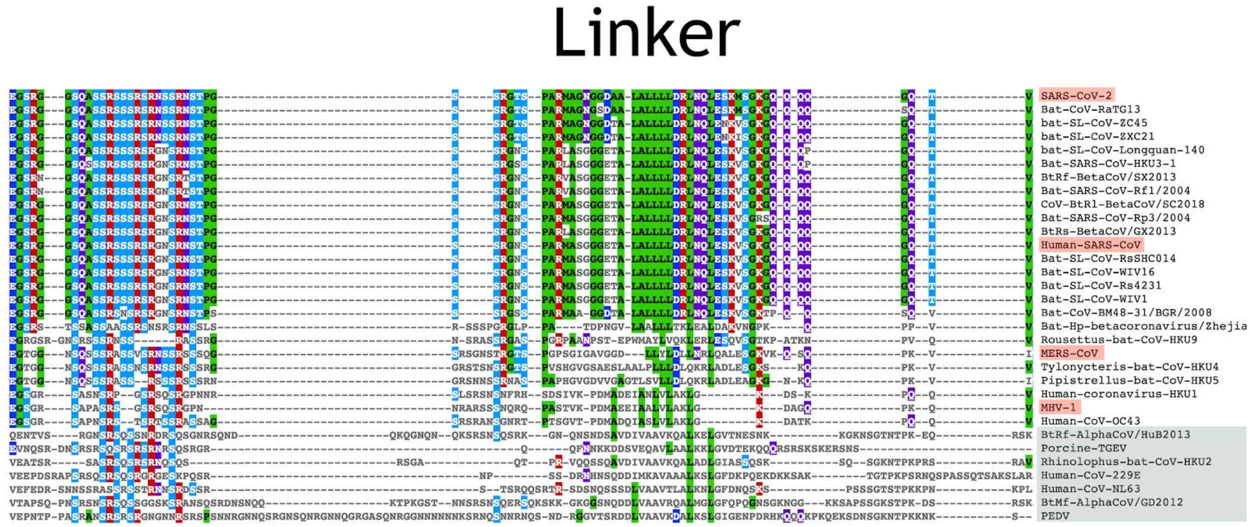


**Fig. S3. Sequence alignment of the coronavirus linker (LINK)**

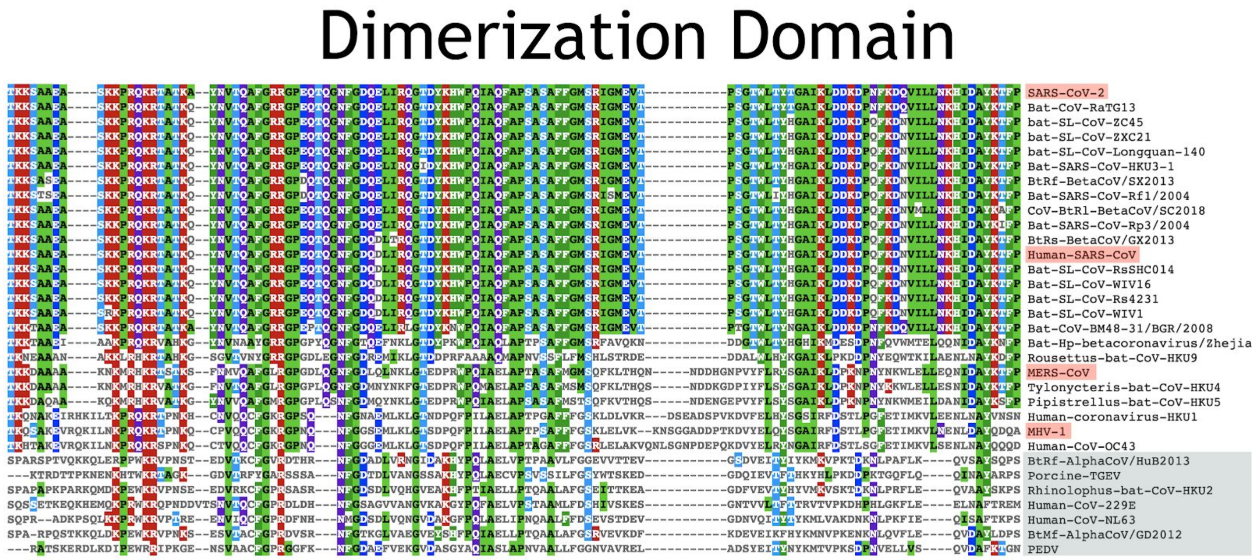


**Fig. S4. Sequence alignment of the coronavirus dimerization domain**

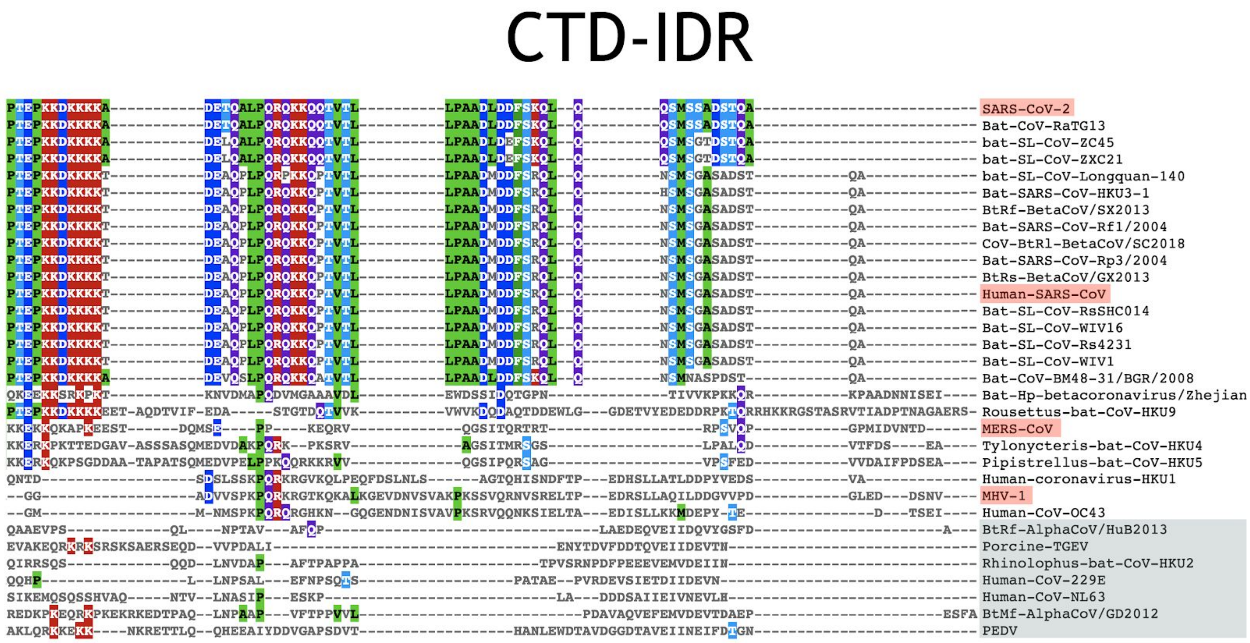


**Fig. S5. Sequence alignment of the coronavirus C-terminal domain (CTD)**

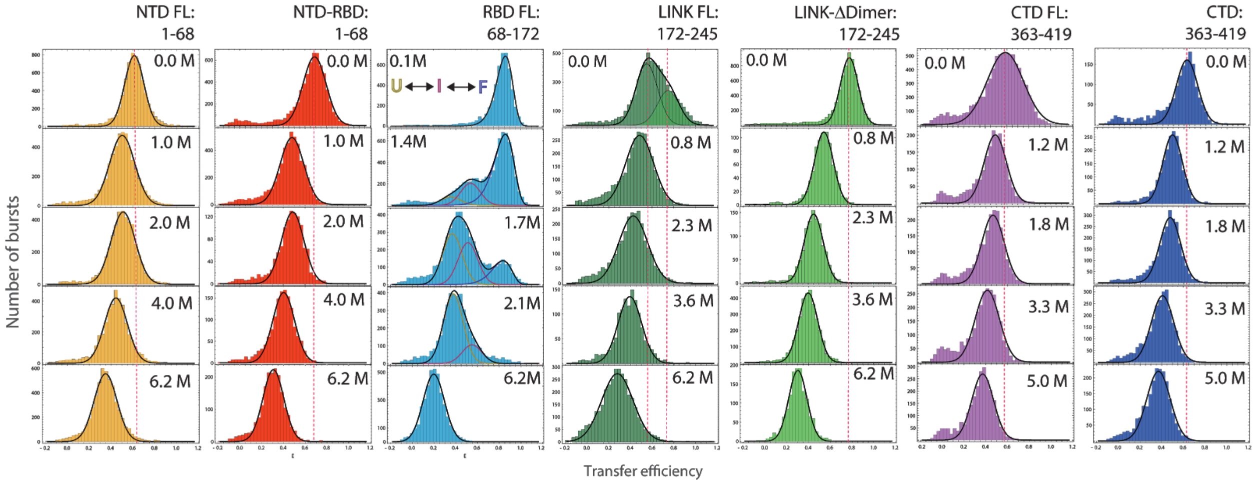


**Fig. S6. Histograms of transfer efficiency distributions across GdmCl concentrations for NTD FL** (orange)**, NTD-RBD** (red)**, RBD FL** (cyan)**, LINK FL** (dark green)**, LINK-ΔDimer** (green)**, CTD FL** (purple) **and CTD** fragment (blue) **constructs.**

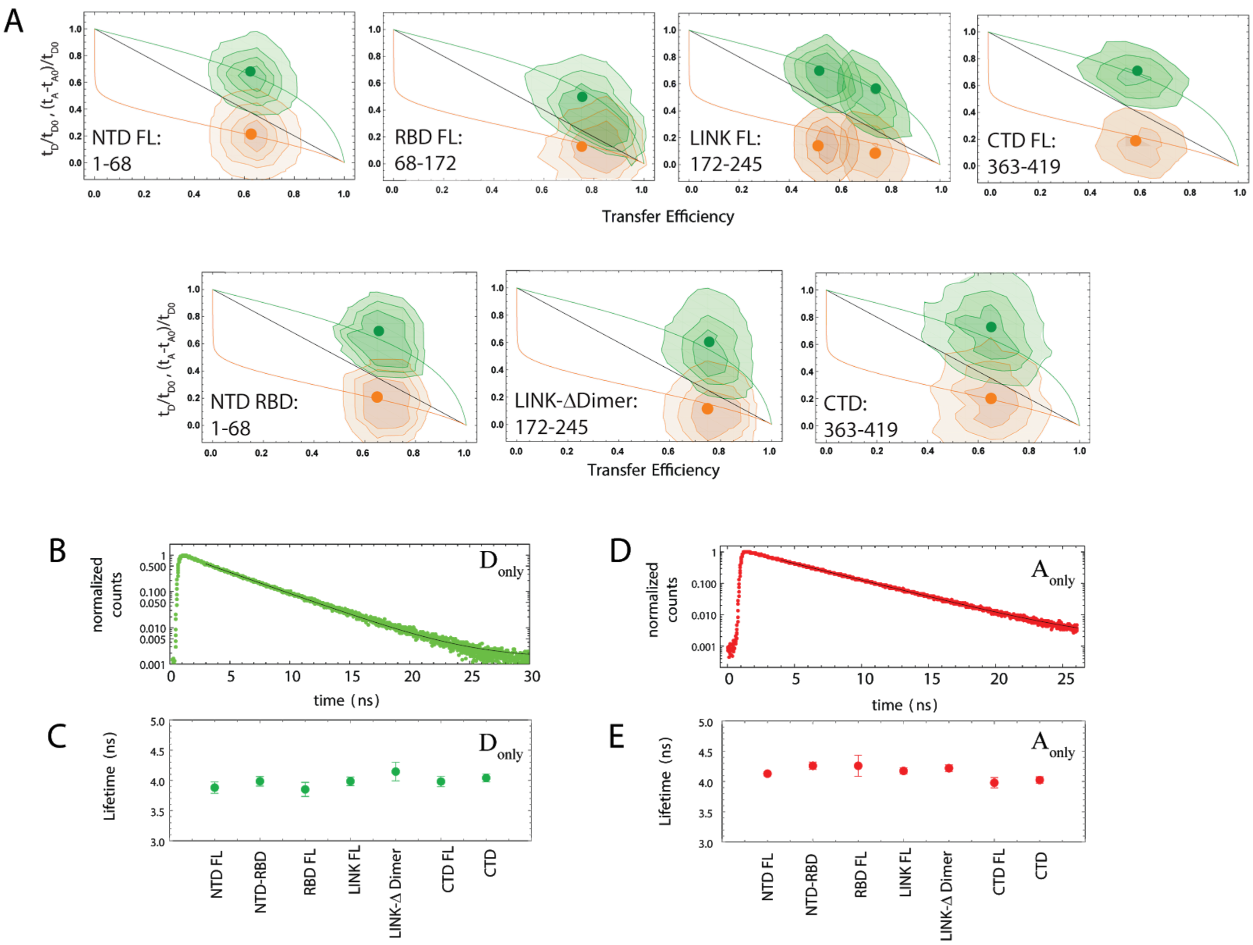


**Fig. S7. Dependence of fluorescence lifetime on transfer efficiency. A.** NTD FL, RBD FL, LINK FL, CTD FL, NTD RBD, LINK-ΔDimer, and CTD construct. Black line: linear dependence expected for a rigid molecule. Green line: the donor lifetime (normalized by the donor lifetime in absence of FRET: $t_D/t_{D0}$) in the limit of dynamics much faster than the burst duration but slower than the fluorophore lifetime. Orange line: the acceptor lifetime delay (normalized by the donor lifetime in absence of FRET: $(t_A\text{-}t_{A0})/t_{D0}$). The green and orange contour plots represent the corresponding distributions of donor lifetime and acceptor lifetime delay as observed in single-molecule experiments under native conditions (**Fig. 2A, 3A, 4A**). The green and orange dots represent the mean value of the measured distributions. **B.** Example of lifetime measurements extracted from the donor-only population and corresponding tail fit. **C**. Observed lifetimes for each construct under aqueous buffer conditions as extracted from the tail fit. **D.** Example of acceptor lifetime measurement from the acceptor only population and corresponding tail fit. **E.** Corresponding acceptor lifetime in aqueous condition for each construct. No significant dynamic quenching is observed in both donor and acceptor. This does not exclude the possible occurrence of static quenching (see **Fig. S12**)

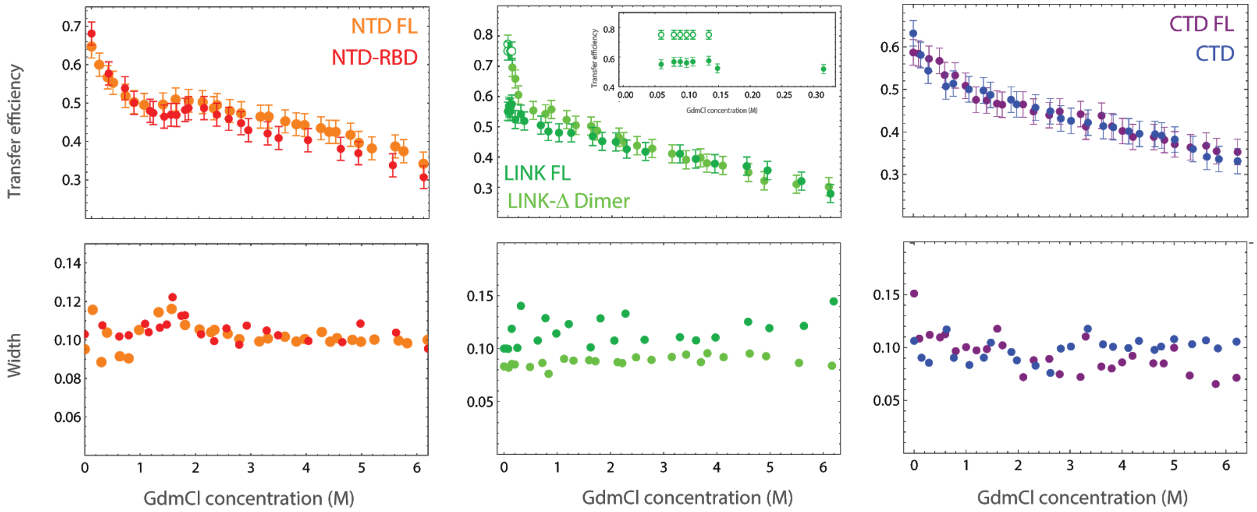


**Fig. S8. Mean transfer efficiency and width of NTD FL vs NTD-RBD, LINK FL vs LINK-ΔDimer, CTD-FL vs CTD fragment across GdmCl concentration.** The mean transfer efficiency of the NTD FL domain (orange) exhibits a plateau between 1 and 2 M GdmCl; at the same concentration we observe a small but systematic increase in the amplitude of the transfer efficiency distribution hinting to the coexistence of two populations in slow exchange with very similar transfer efficiencies. The same behavior is closely reproduced by the truncated variant NTD-RBD (red). The LINK FL (dark green) exhibits two distinct populations at very low GdmCl concentration (open and close circles), suggesting a strong contribution of electrostatics in favoring one of the two configurations. Inset shows coexistence of the two states between 0 M and 0.75 M GdmCl. The truncated variant LINK-ΔDimer (green) shows a continuous collapse that interpolates the two positions observed for the LINK FL, suggesting interaction of the LINK with itself or with the RBD domain in absence of the DIMER domain. Finally, the CTD FL (blue) and the CTD fragment (purple) exhibit similar conformations across denaturant concentrations. The small increase in the width of the transfer efficiency distribution that may reflect the formation of local structure under native conditions (e.g. the putative helical binding motif).

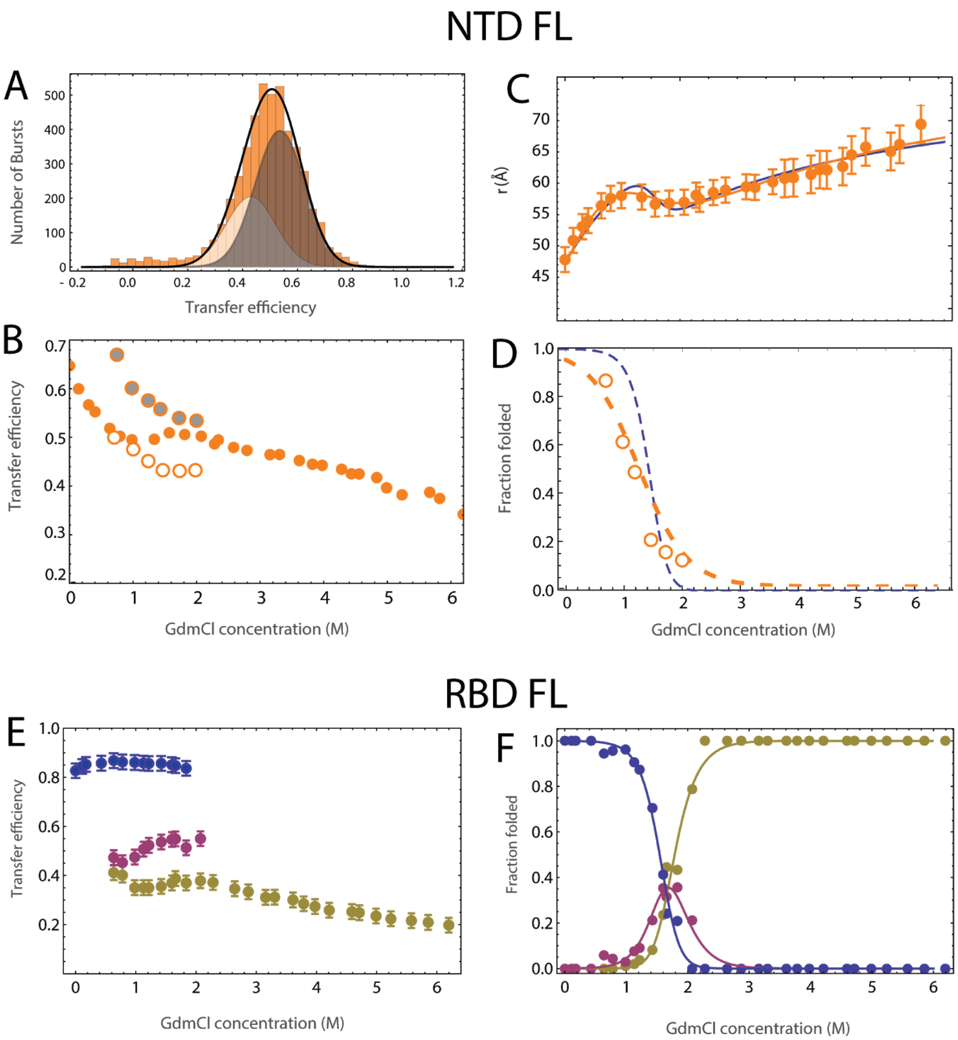


**Fig. S9. Fit of NTD construct with two populations compared to folding of RBD domain.** To address the change in amplitude that occurs from the NTD construct between 1 and 2 M GdmCl, we attempt a fit of the NTD FL data using two populations with a fixed width equal to average width below 1 M and above 2 M GdmCl (see for comparison **Fig. S8**) **A.** Fit of the transfer efficiency histogram at 1.5 M GdmCl. The white- and gray- shaded areas reflect fits to the "folded RBD" population and to the "unfolded RBD" population. *Central panel:* Comparison of transfer efficiencies with a single fit (solid orange circles, compare **Fig. S8**) and from the two populations: gray solid circles for the "unfolded RBD" population and unfilled circles for the "folded RBD" population. *Lower panel*: Fraction folded estimated from the fit with **Eq. S7** compared to the fraction of "folded RBD" obtained from computing the ratio between the area under "folded RBD" species and the total histogram area.

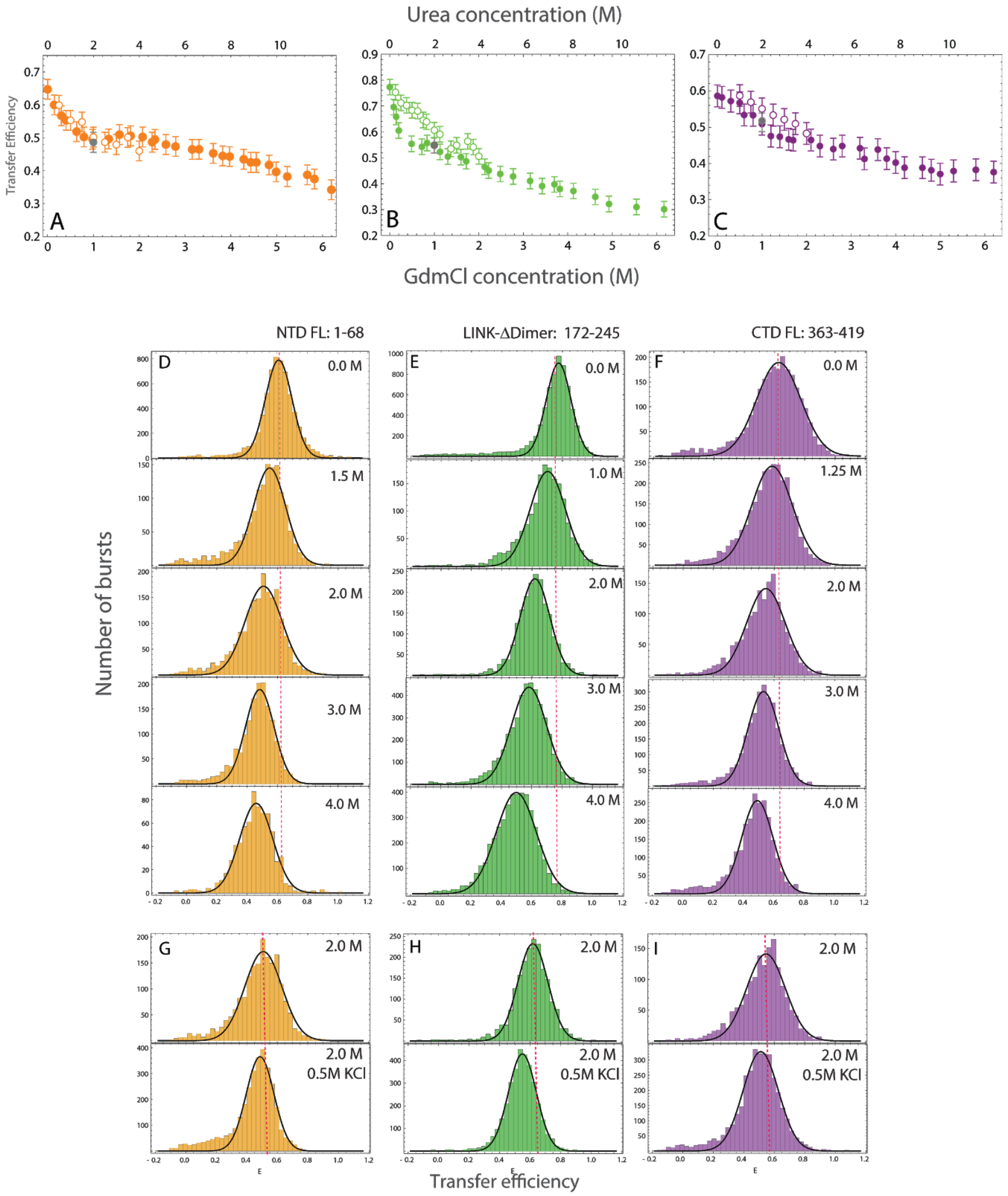


**Fig. S10. Effects of Urea denaturation on NTD FL, LINK-ΔDimer, and CTD FL. A-C**. Comparison of Urea (open circles) and GdmCl (close circles) effects on the transfer efficiencies of NTD FL (orange), LINK-ΔDimer (green), and CTD-FL (purple). The Urea range is rescaled by a factor of 2 compared to the GdmCl range to account for the different denaturing effect. Grey dots correspond to the same concentration of Urea with the addition of 0.5 M KCl. **D-F.** Examples of transfer efficiencies distribution as function of Urea. **G-I.** Comparison between 2 M Urea histograms in presence and absence of 0.5 M KCl.

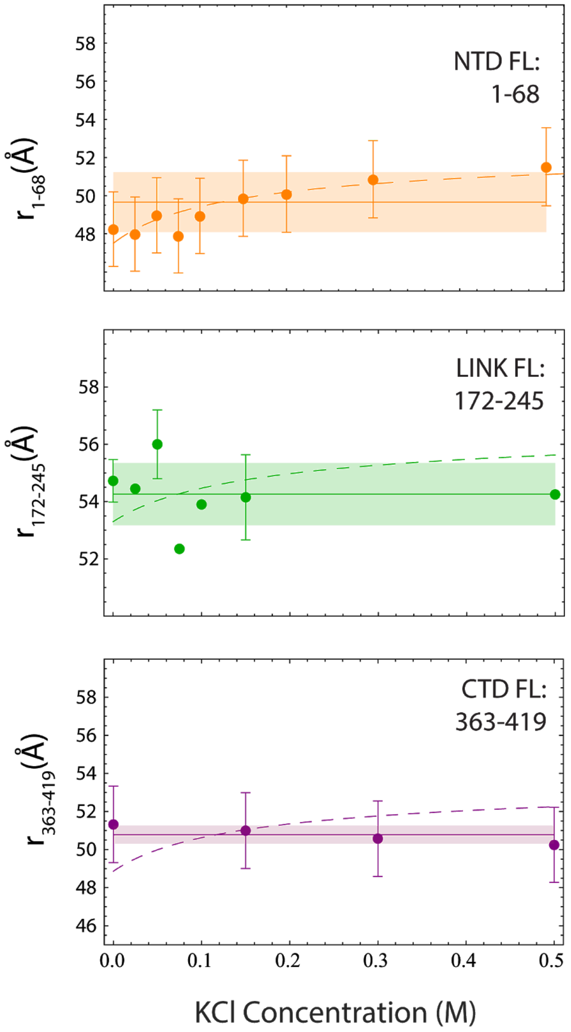


**Fig. S11. Interdye distances of NTD, LINK, CTD in presence of salt (KCl).** *Upper panel*: root mean square interdye distance between position 1 and 68. Dashed line: fit according to the Higgs & Joanny model **(Eq. S11-12)** predicts a comparable change to the one observed. *Central panel:* root mean square interdye distance between position 172 and 245. Dashed line: fit according to the Higgs & Joanny model (**Eq. S11-12**) predicts a comparable change to the one observed. Solid line and shaded area: average value of the root-mean-square interdye distance across all salt conditions and corresponding standard deviation. The standard deviation is comparable to the measurement error. *Lower panel*: root mean square interdye distance between position 363 and 419. Dashed line: fit according to the Higgs & Joanny model (**Eq. S11-12**) does not capture the observed trend. This can be possibly explained considering the significant predicted population of helical conformations in the CTD. Solid line and shaded area: average value of the root-mean-square interdye distance across all salt conditions and corresponding standard deviation.

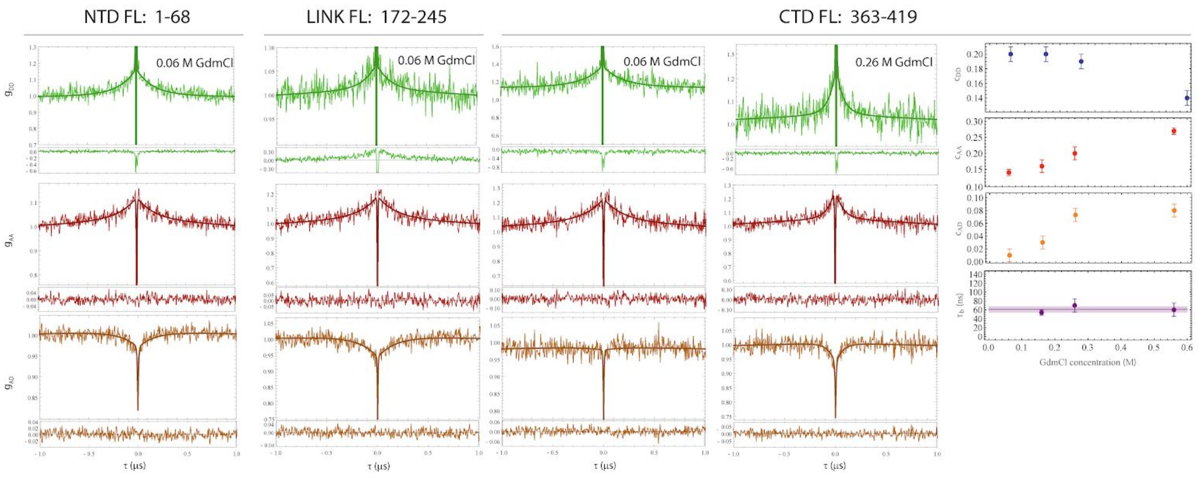


**Fig. S12. Chain dynamics measured via ns-FCS.** Nanosecond FCS measurements for the NTD, LINK, and CTD constructs provide a measure of the dynamics on the nanosecond timescale. All correlations are normalized to the value measured at 1 μs for highlighting the amplitude relative to the reconfiguration term. The donor-donor (green), acceptor-acceptor (red), and donor-acceptor (orange) correlation are fitted to a global model that accounts for antibunching, FRET dynamic populations, and triplet. The acceptor-donor correlation shows a clear anticorrelated change for NTD FL and LINK FL in the signal that reflects the anticorrelated nature of the donor-acceptor energy transfer as a function of distance: an increase in acceptor reflects a decrease in donor. The CTD FL cross-correlation exhibits a flat behavior, which is consistent with absence of dynamics or compensation between two populations, one anti-correlated (dynamic) and one correlated (static). The observation of a correlation decay in the donor-donor and acceptor-acceptor autocorrelations, with characteristic times of 80 ± 20 ns and 110 ± 20 ns respectively, suggests a contribution of static quenching by Photo-induced Electron Transfer (PET) [58] [34]. Addition of GdmCl (e.g., 0.26 M) causes a decrease in the transfer efficiency distribution width **(Fig. S8)** and leads to the appearance of an anticorrelated increase in the cross-correlation of CTD. A plot of the corresponding change in amplitude for the donor-donor, acceptor-acceptor, and acceptor-donor correlation is shown for comparison. We interpret the decrease in the donor-donor correlation and the increase in the acceptor-acceptor and acceptor-donor correlations as the result of destabilization of the quenched species in favor of the dynamic population. A decay correlation time can be globally fitted starting from 0.16 M GdmCl and appears to be constant across the measured values, up to 0.6 M GdmCl. The average decorrelation time $t_{CD}$ is equal to 61 ± 7 ns.

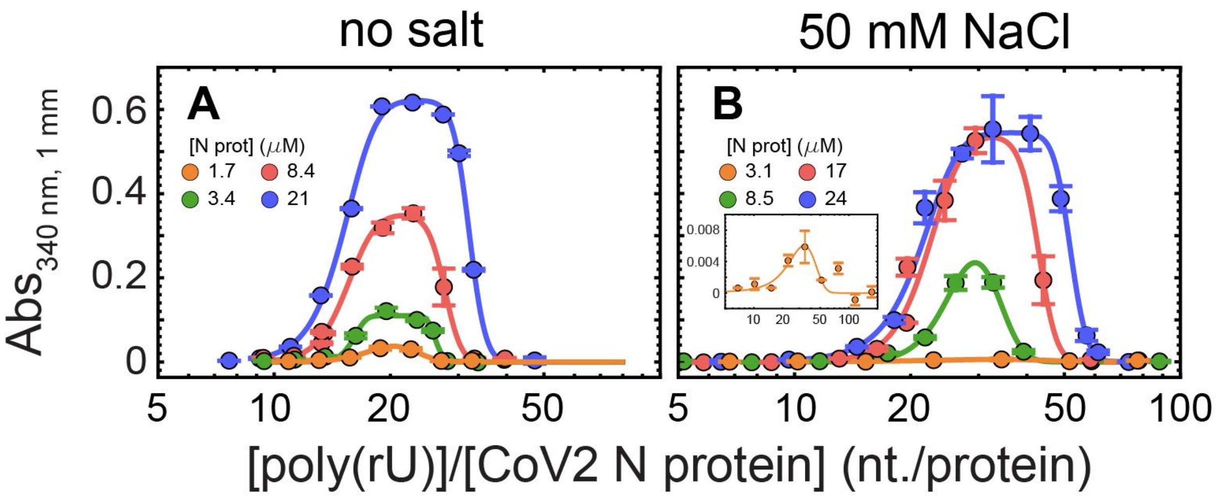


**Fig. S13. Turbidity experiments plotted against RNA/protein ratio.** Representative turbidity titrations with poly(rU) in 50 mM Tris, pH 7.5 (HCl) at room temperature, in absence of added salt (A) and in presence of 50 mM NaCl (B), at the indicated concentrations of N protein. On the x-axis, the concentration of poly(rU) is rescaled for the protein concentration. Points and error bars represent the mean and standard deviation of 2-4 consecutive measurements from the same sample. Solid lines are simulations of an empirical equation fitted individually to each titration curve. An inset is provided for the titration at 3.1 µM N protein in 50 mM NaCl to show the small yet detectable change in turbidity on a different scale. Interestingly, within the experimental error, we observe a clear alignment of the turbidity curves with a maximum at ~20 nucleotides per protein in the absence of added salt (A) and ~30 nucleotides per protein in the presence of 50 mM NaCl (B).

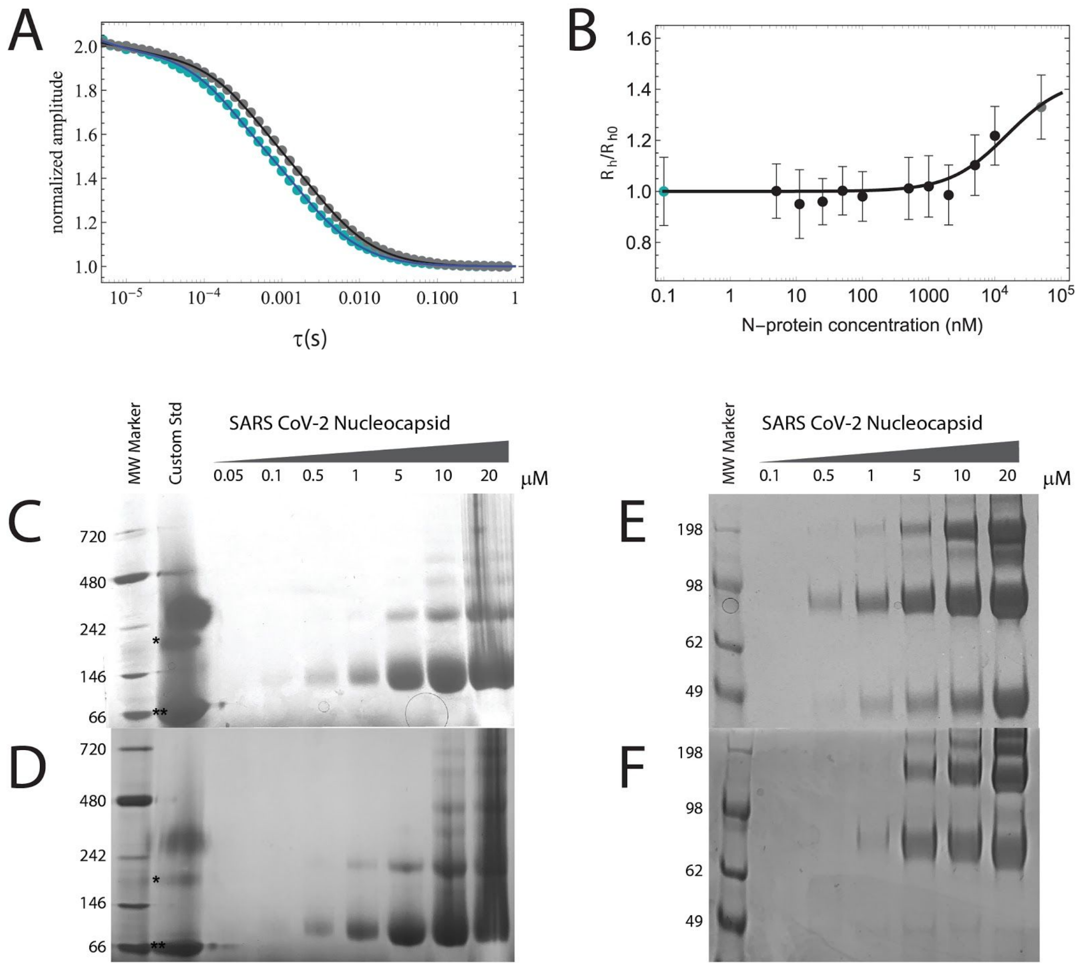


**Fig. S14. Testing SARS-CoV-2 N protein oligomerization. (A-B)** Fluorescence Correlation Spectroscopy (FCS) of full-length SARS-CoV-2 N protein as a function of protein concentration. (A) FCS traces of 100pM Alexa 488/Alexa 594 N protein labeled at positions 363 and 419 in the absence (blue dots) and the presence (gray dots) of 50 μM unlabeled N protein. (B) Hydrodynamic radius of SARS-CoV-2 N protein obtained from FCS trace analysis (blue dot: 100pM labeled N protein; gray dot: 100pM labeled N protein + 50μM unlabeled N protein). **(C-D)** NativePAGE of full-length SARS-CoV-2 N protein in 20 mM NaPi pH 7.4 as a function of protein concentration in the presence of 200 mM NaCl (C) and in the absence of added salt (D). 'Custom Std' lane contains Alcohol Dehydrogenase ( * , 150 kDa) and Bovine Serum Albumin (** , 66 kDa). (E-F) SDS PAGE of crosslinked full-length SARS-CoV-2 N protein in 20 mM NaPi pH 7.4, 1.25 mM DSS as a function of protein concentration in the presence of 200 mM NaCl (E) and in the absence of added salt (F).

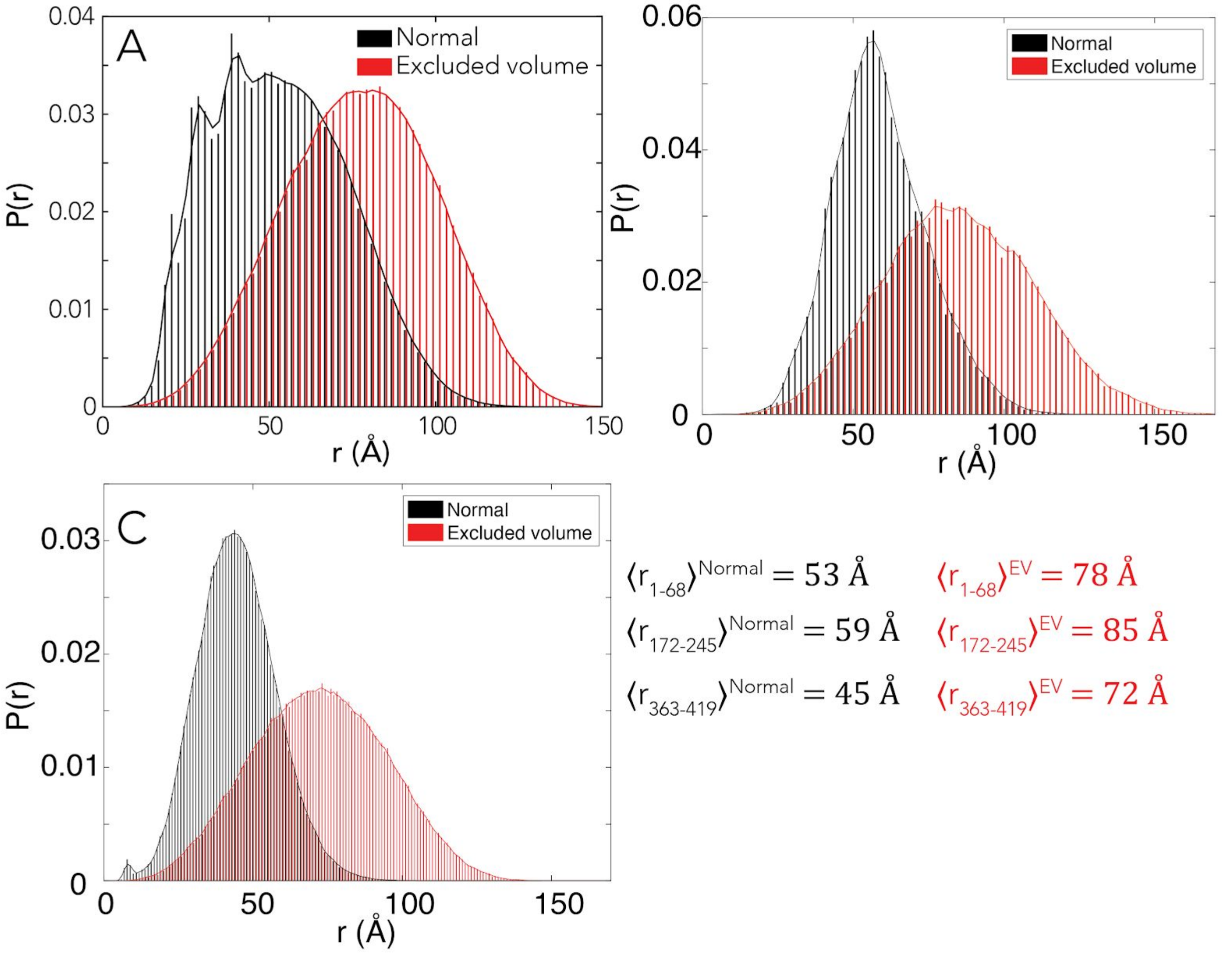


**Fig. S15. Distributions of inter-residue distance from ABSINTH simulations (black) vs. excluded volume simulations (red).** Comparison of simulations with the full ABSINTH Hamiltonian ('normal', black) against simulations performed in the excluded volume (EV, red) limit for **(A)** NTD in the NTD-RBD context, **(B)** LINK in the NTD-LINK-DIM context, and **(C)** CTD in the DIM-CTD context. In all three cases the EV simulations are performed in the analogous structural contex, and report substantially larger average distances than the ABSINTH simulations, as expected given the absence of any attractive intramolecular interactions. The distances reported from the EV simulations are also slightly more expanded than under fully denatured conditions, consistent with systems studied previously (see [7,59]).

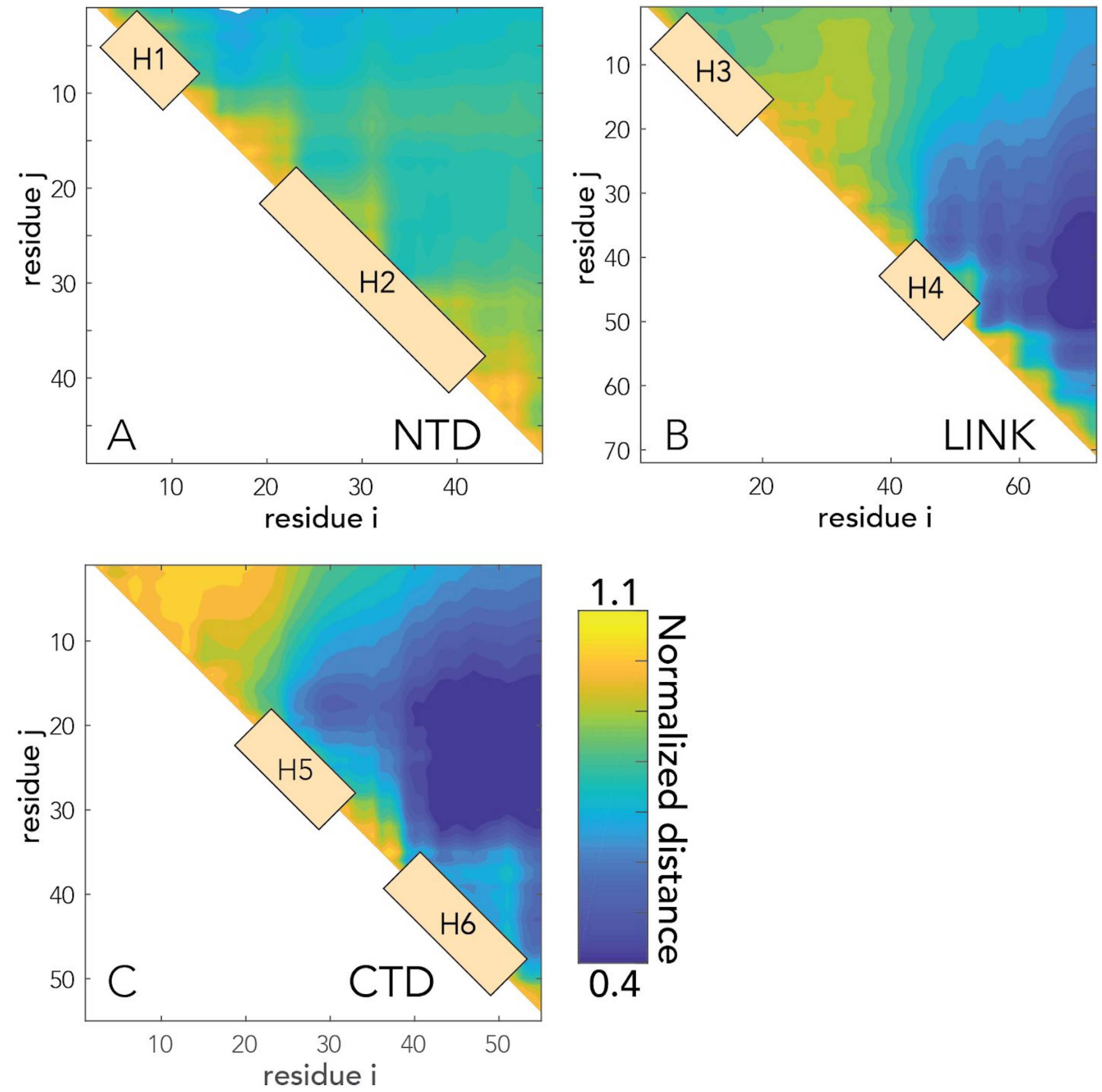


**Fig. S16. Scaling maps for IDR-only simulations.** Scaling maps report on the normalized distance between pairs of residues, where normalization is done by the distance expected if the IDRs behaved as self-avoiding chains in the excluded-volume limit. Scaling maps for IDR-only simulations of the **(A)** NTD, **(B)** LINK and **(C)** CTD. For each sequence, transient helices are annotated on the scaling maps. Note that in the LINK we observe interaction between the C-terminal region of the LINK and H4, while H3 does not interact with any parts of the sequence. Similarly, in CTD we see extensive intramolecular interactions between H5 and H6.

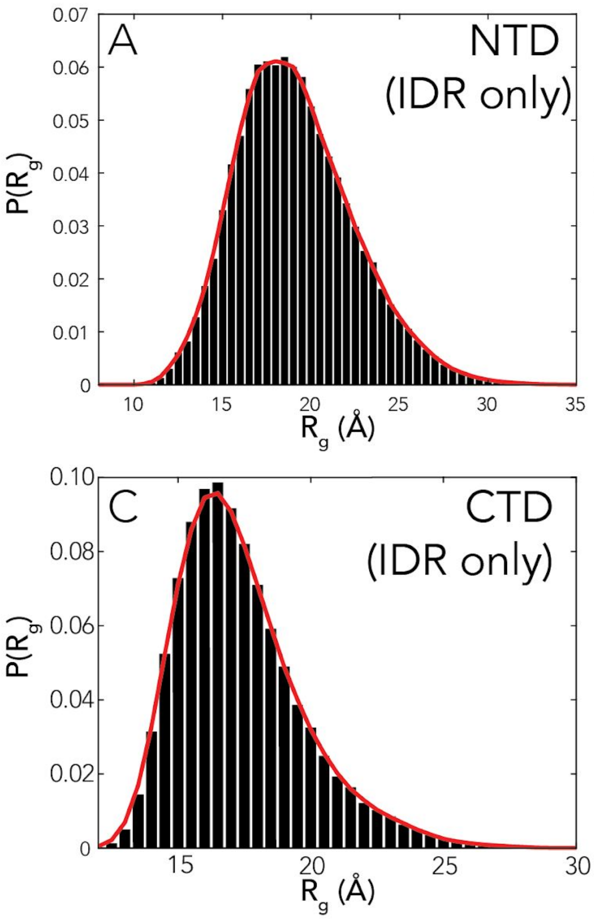


**Fig. S17. Distributions for the radius of gyration ($R_g$) of for IDR-only simulations.** $R_g$ distributions for **(A)** NTD, **(B)** LINK and **(C)** CTD. Average $R_g$ for each IDR in isolation is 19.1 Å (NTD), 21.4 Å (LINK), and 17.1 Å (CTD).

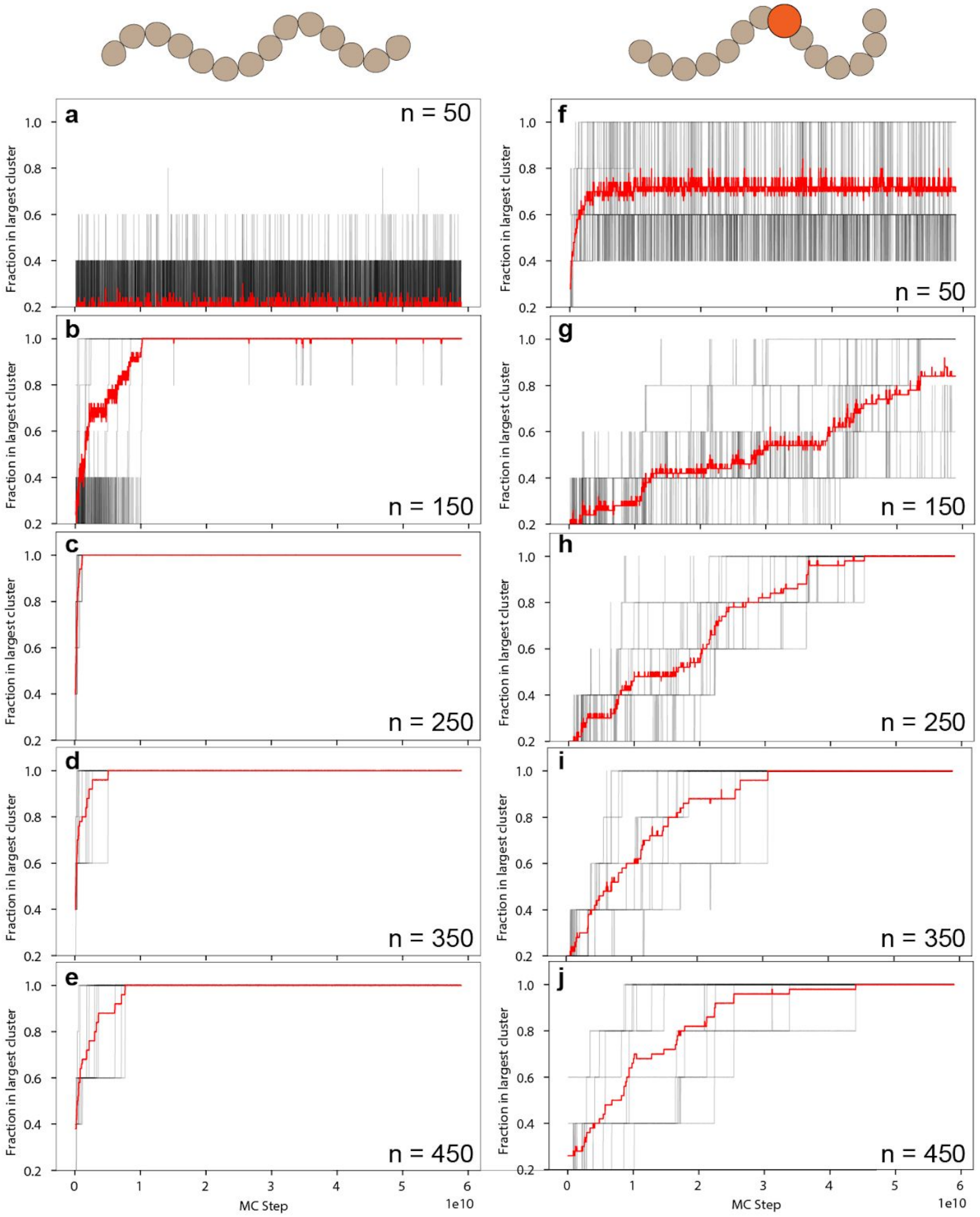


**Fig. S18. Kinetic Monte Carlo simulations reveal slow kinetics of condensate fusion.**
Our simulations in **Fig. 6** reveal single-polymer condensates in the presence of a high-affinity binding site, whereas multi-chain droplets assemble in the absence of a high-affinity binding site. To further explore the origin of single-polymer condensates we ran extensive Monte Carlo simulations using an approximate kinetics scheme (that includes cluster translation moves) to examine the pseudo-kinetics of assembly. Black lines in each panel correspond to individual simulation trajectories, while red lines report on the average behavior over ten independent simulations. *n* reflects the number of binder chains in each simulation. To assess the kinetics of

assembly, we asked what fraction of the total number of polymers are found in the largest cluster. Under conditions in which a single droplet forms 100% of the polymer chains will be found in the largest cluster. Panels a-e report on behavior for polymers without a high affinity binding site. In all cases within $10^9$ Monte Carlo steps every independent simulation has converged on a single multichain droplet that represents the thermodynamic minimum expected for a two-phase equilibrium. Panels f-j report on identical simulations performed with a high affinity binding site. While these simulations trend towards or reach a single multichain condensate, the presence of a high-affinity binding site substantially retards the assembly kinetics, revealing a large regime over which single-polymer condensates are metastable.

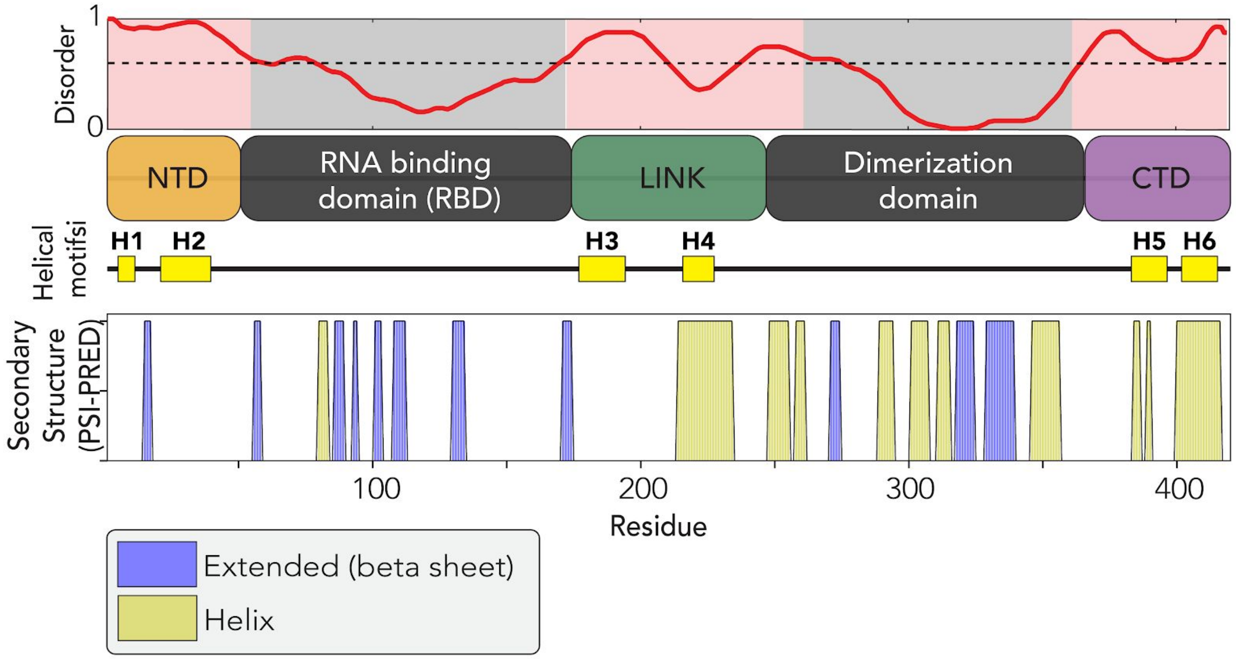


**Fig. S19. Comparison of secondary structure in IDRs from bioinformatics predictions**. We computed secondary structure propensities for the full-length protein using the PSI-PRED prediction server [60]. This analysis correctly identifies helices H4, H5 and H6, but fails to identify those H1, H2 and H3. Helix H3, H4 and H6 have been similarly identified by NMR and/or hydrogen-deuterium exchange mass spectroscopy [16,61,62]. These results demonstrate that our simulations are able to identify predicted helices but, furthermore, find helices that conventional structural bioinformatics software fails to correctly identify.
